# Insights into cargo sorting by SNX32 in neuronal and non-neuronal cells: physiological implications in neurite outgrowth

**DOI:** 10.1101/2022.11.04.515170

**Authors:** Jini Sugatha, Amulya Priya, Prateek Raj, Ebsy Jaimon, Anju Jose, Sunando Datta

## Abstract

Sorting nexins (SNX) are a family of proteins containing the Phox homology domain, which shows a preferential endo-membrane association and regulates cargo sorting processes. Even with the vast amount of information unveiled systematically, the underlying mechanism of sorting remains elusive. Here, we established that SNX32, a SNX-BAR (Bin/Amphiphysin/Rvs) sub-family member, is associated with SNX4 via its BAR domain. We identified A226, Q259, E256, R366 of SNX32, and Y258, S448 of SNX4 at the interface of these two SNX proteins that are important for maintaining the association. Via its PX domain, SNX32 interacts with the Transferrin receptor (TfR) and Cation Independent Mannose-6-Phosphate Receptor (CIMPR). We showed that the conserved F131 in its PX domain is important in stabilising the above interactions. Silencing of SNX32 led to a defect in intracellular trafficking of TfR and CIMPR, which could be rescued by overexpressing shRNA-resistant snx32. We also showed that both individual domains play an essential role in trafficking. Our results indicate that SNX4, SNX32 and Rab11 may participate in a common pathway regulating transferrin trafficking; however, the existence of an independent pathway for Rab11 and SNX32 could not be completely ruled out. Further, we established that the PX domain of SNX32 could bind to PI(3)P and PI(4)P, suggesting a possible explanation for its sub-cellular localization. Taken together, our study showed that SNX32 mediate the trafficking of specific cargo molecules along distinct pathway via its PX domain-directed binding to phosphoinositides and its BAR domain-mediated association with other SNX family members. Further, using SILAC-based differential proteomics of the wild type and the mutant SNX32, impaired in cargo binding, we identified Basigin (BSG), an immunoglobulin super family member, as a potential interactor of SNX32 in SHSY5Y cells. We then demonstrated that SNX32 binds to BSG through its PX domain and facilitates its trafficking to the cell surface. In Neuro-Glial cell lines, the silencing of SNX32 led to defects in neuronal differentiation. Moreover, abrogation in lactate transport in the SNX32 depleted cells led us to propose that the SNX may contribute to maintaining the neuro-glial coordination via its role in BSG trafficking and the associated Monocarboxylate transporter activity.

## INTRODUCTION

Approximately 25% of the human genome encodes for integral membrane proteins (around 5,500 proteins). Efficient intracellular transport of these proteins and associated proteins and lipids (together termed ‘cargos’) is essential for organelle biogenesis, maintenance, and quality control. This is an extensive field that encompasses the secretory, endosomal, lysosomal, and autophagic pathways, all fundamental features of eukaryotic cells. Efficient integration of these pathways is essential for cellular organization and function, with errors leading to numerous diseases, including those associated with ageing and neurodegeneration.

The endosomal network is composed of a series of distinct compartments that function to efficiently sort and transport cargo proteins between two distinct fates: either sorting for degradation in the lysosome or retrieval from this fate for recycling and transport to an array of organelles that include the cell surface, and the biosynthetic and autophagic pathway. The sorting nexin family is a conserved group of proteins with diverse roles in regulating the function of the endosomal network. Other than the PX-driven lipid specificity, much of the functional diversity displayed by the family could be apprehended by its ability to indulge in protein dimerization and oligomerizations^1^. It is interesting to note that individual SNX-BAR proteins incapable of remodelling membrane *in vitro* (SNX5, SNX6, SNX7, SNX30 and SNX32 etc.) are observed to undergo heterodimeric interactions with SNX-BAR proteins possessing an intrinsic membrane remodelling capacity (SNX1, SNX2, SNX4, SNX8 etc.)^2^. Protein-protein interactions thus add layers of intricate regulations, increasing the complexity of the system^2^.

Conserved throughout eukaryotes, defects in the function of these proteins are linked with various pathologies, including Alzheimer’s disease (AD)^3^, where they are associated with trafficking and processing^1, 4–6^, Down’s syndrome (DS)^7^, cancer^8^, schizophrenia, hypertension^9^, thyroid disorders, and epilepsy^7^. The pathological implications of sorting nexin is an emerging area of extensive investigation^7^.

Sorting nexin 32 (SNX32), also known as SNX6b^10^, on account of sequence similarity to its paralogue sorting nexin-6^6^, is an underexplored member of the sorting nexin-containing BAR domain (SNX-BAR) sub-family of sorting nexins^6^. SNX32 is a component of the recently identified Endosomal SNX–BAR sorting complex for promoting exit 1 (ESCPE-1), a heterodimer of SNX1/SNX2 associated with SNX5/SNX6/SNX32. ESCPE-1 regulates the sequence-dependent sorting of transmembrane cargo by recognizing and binding to a specific bipartite motif present within their cytosolic tail, an interaction mediated by the SNX5, SNX6 and SNX32 subunits^1, 10–12^. The previous reports suggest that the ESCPE-1 complex identifies a bipartite signal sequence (ΦxΩxΦ where Φ is hydrophobic and Ω is aminoacid with aromatic side chains) present in the cytosolic tail of the prototypical cargoes and assists in its retrieval and recycling to TGN as well as to the surface^1^. Even though it is referred that similar to its homologues SNX5/SNX6, SNX32 contributes to the trafficking of the CIMPR, it is yet to be demonstrated experimentally. Moreover, SNX32 is absent in non-higher metazoans, and in humans, its expression is skewed to the brain, making it a compelling sorting nexin to understand within the context of neurobiology (http://www.gtexportal.org/home/gene/SNX32). In spite of heavy reliance on endocytic trafficking for functional fluidity in neurons, the underlying interlaced network of sorting leading to recycling or degradation remains elusive^13, 14^. In this study, we have investigated the interaction of SNX32 within the SNX family, its lipid affinity, and its role in the intracellular trafficking of CIMPR, Transferrin receptor and Basigin. We begin to define the function of SNX32 in neuronal endosomal sorting, revealing an important contribution to the process of neurite outgrowth.

## RESULTS AND DISCUSSION

### SNX32 UNDERGOES BAR DOMAIN-ASSISTED INTERACTION WITH SNX1 AND SNX4

The characteristic ability of the SNX-BAR family of proteins to undergo oligomerization directed us to examine the ability of SNX32 to participate in such interactions within the family. We utilized a GFP nanobody-based immunoprecipitation (GBP) assay for the same. The GST-GBP immunoprecipitation assay was carried out by immobilizing GST-GBP on GST beads and incubating with HEK293T cell extracts overexpressing GFP/GFP-SNX1/GFP-SNX4/GFP-SNX8/GFP-SNX32 and HA-SNX32. The eluates were resolved through SDS-PAGE and subjected to immunoblotting using an anti-GFP and anti-HA antibody. Immunoblotting using an anti-HA antibody showed that GFP-SNX1(0.98), GFP-SNX4(0.92), GFP-SNX8(0.62) and SNX32(0.85) were efficiently precipitating HA-SNX32(Fig.1A), which is also in agreement with the report by Sierecki, E. *et al.* ^15^. To identify the limiting region on SNX32 that contributes to these interactions, we generated two deletion constructs, an SNX32ΔN, spanning the BAR domain15 (167-404) and SNX32ΔC, spanning the PX domain^16^ (1-166) (Fig.S1A). We performed a GFP nanobody-based immunoprecipitation (GBP) assay where GST-GBP was immobilized on GST beads and incubated with HEK293T cell lysate overexpressing GFP-SNX1/GFP-SNX4 and HA-SNX32FL/HA-SNX32ΔC/HA-SNX32ΔN. The eluates were resolved through SDS-PAGE and subjected to immunoblotting using an anti-GFP, anti-HA antibody. We found that GFP-SNX4 (Fig. 1B)/GFP-SNX1 (Fig.S1B) precipitated HA-SNX32FL and HA-SNX32ΔN but not HA-SNX32ΔC.To assess whether the two SNXs are involved in direct interaction, we attempted to purify individual interacting partners from bacterial expression systems. Though His-SNX32ΔC was purifiable (Fig S1C), the full length, SNX32ΔN as well as the GST-SNX1/His-SNX32ΔN comple[, were unavailing due to the insolubility of the expressed proteins (Fig. S1 D-E).

**Figure 1.**
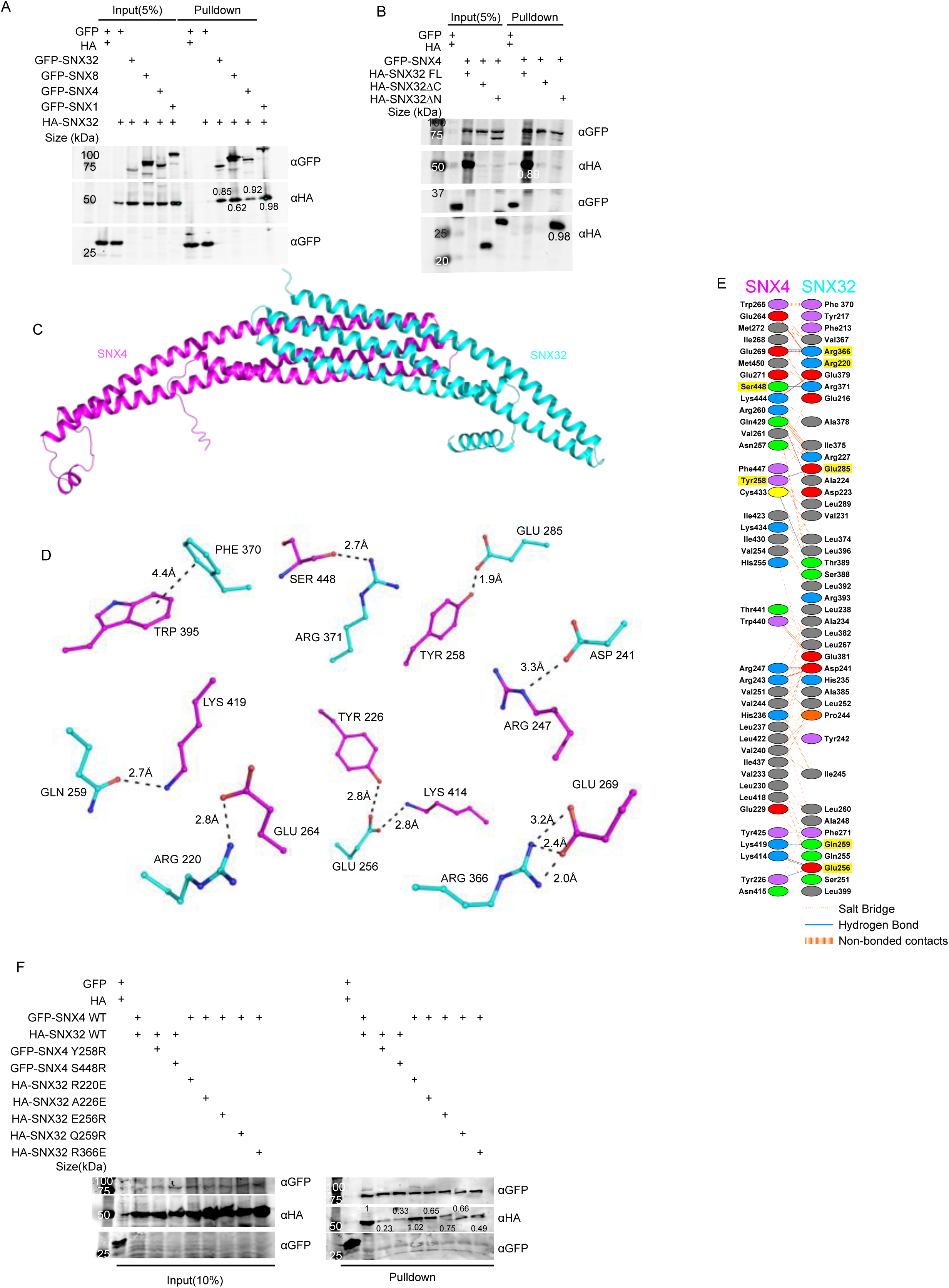
SNX32 undergoes BAR domain-mediated association with SNX4. A) GBP co-immunoprecipitation of GFP /HA-tagged SNX–proteins transiently transfected in HEK293T cells showing GFP-SNX1/ GFP-SNX4/ GFP-SNX8 or GFP-SNX32 efficiently precipitating HA-SNX32 (representative immunoblot out of 3 biological replicates, values represent the ratio of HA to GFP band intensity). B) Co-immunoprecipitation of GFP/HA-tagged SNX–proteins transiently transfected in HEK293T cells showing GFP-SNX4 precipitating HA-SNX32ΔN (representative immunoblot out of 3 biological replicates, values represent the ratio of HA to GFP band intensity). C)homology model of BAR domains of SNX32-SNX4 complex. D) Schematic depiction of the amino acid residues lining the heterodimeric interface of SNX32(cyan)-SNX4(magenta). E) Polar interactions present at the dimeric interface of SNX32(cyan)-SNX4(magenta). F) GBP co-immunoprecipitation of GFP/HA-tagged SNX– proteins transiently transfected in HeLa cells showing difference in the amount of GFP-SNX4 precipitated HA-SNX32 mutants (representative immunoblot out of 3 biological replicates, values represent the ratio of HA to GFP band intensity).

### SNX32 AND SNX4 INTERACTION IS MEDIATED BY THE RESIDUES IN THE BAR DOMAIN INTERFACE OF THE SNXS

Our experimental data showed that HA-SNX32 could interact with GFP-SNX1/GFP-SNX4 through its BAR domain. In order to identify the critical residues in maintaining the association between these SNXs, we predicted the structure of BAR domain heterodimers using the Alphafold2-Multimer (ColabFold) program ^17–19^.

Generated models showed a high overall predicted Local Distance Difference Test (pLDDT) score, suggesting the high accuracy of the models. A few stretches (residues at the kinks between two helices) and several terminal residues have low pLDDT scores. However, most of the dimeric interface regions have a pLDDT score of more than 85. To assess the accuracy of our predictions, we modelled the homodimer of the SNX1-BAR domain for which the X-ray crystal structure (PDB ID: 4FZS) as well the cryo-EM structures (PDB ID: 7D6D and 7D6E) was already available. A comparison of the predicted model with the experimentally known SNX1-BAR homodimer revealed that the predicted model is highly consistent with the cryo-EM structure, thus resembling the membrane-bound SNX1 homodimer.

The predicted heterodimeric models of SNX32-BAR with SNX1-BAR and SNX4-BAR adopt a banana-shaped structure, a characteristic of conventional BAR domains (Fig. 1 C-E, Fig. S1 F-K). In the case of the SNX1-SNX32 BAR heterodimer (Fig. S1 F-H), the buried surface area (BSA) of SNX1-BAR and SNX32-BAR is 2572 Å^2^ and 2519 Å^2^, respectively. Similarly, the BSA of SNX4-BAR and SNX32-BAR in the SNX4-SNX32 BAR heterodimer (Fig. 1 C-E) is 2514 Å^2^ and 2505 Å^2^, respectively. High BSA represents the formation of strong heterodimers. On a closer look at the interphase of the SNX4-SNX32 complex, there are 7 salt bridges and 9 hydrogen bonds, whereas, in the case of the SNX1-SNX32 complex, there are 4 salt bridges and 6 hydrogen bonds. The overlay of SNX1-SNX32 and SNX4-SNX32 revealed that the curvature attained in either case is comparable. The comparison of SNX1-SNX32(Fig. S1 F-H) with SNX32-SNX32 (Fig. S1 I-K) revealed that Y242, R366, and K400 (Fig. S1 G, J) of SNX32 participate in stabilizing both homodimeric and the heterodimeric interface, whereas D223, L399, E402 (Fig. S1 G, H) uniquely mediates the interactions at the heterodimeric interface. Similarly, in the case of SNX4-SNX32, R220, E256 and R366 of SNX32 (Fig. 1 D) participates in both hetero/homodimeric interfaces, whereas D241, Q259, E285, F370, R371(Fig. 1 D, E) participates only in the heterodimeric interface. Of note, F370 is involved in mediating a pi-pi interaction with F264 of SNX32 and W265 of SNX4(Fig. 1 D, E). Overall, the heterodimeric interface of SNX1-SNX32 and SNX4-SNX32 displays extensive hydrophobic interactions along with several polar interactions.

Based on the above model, we next assessed if the interaction between HA-SNX32 and GFP-SNX4 could be disrupted by mutating the amino acids engaged in interactions at the interphase. Single mutations of S448R and Y258E in the GFP-SNX4 BAR domain and single mutations of R220E, E256R, Q259R and R366E in the HA-SNX32 were carried out utilizing site-directed mutagenesis. Additionally, it was previously reported that SNX5S226E mutation disrupts its heterodimeric interaction with SNX1, whereas SNX5S226A doesn’t ^1, 20^; based on that, we looked into the structure of SNX32, which had an Alanine in the 226^th^ position and decided to include A226E in our study.

We performed a GBP-based immunoprecipitation assay where GST-GBP was immobilized on GST beads and incubated with HeLa cell lysate overexpressing GFP-SNX4 WT/ S448R/Y258E and HA-SNX32 WT or GFP-SNX4 WT and HA-SNX32 WT/ R220E/ A226E/ E256R/ Q259R/ R366E. The eluates were resolved through SDS-PAGE and subjected to immunoblotting using an anti-GFP, anti-HA antibody. We found that HA-SNX32 FL was efficiently precipitated by GFP-SNX4 WT but not GFP-SNX4 S448R (0.23) /Y258E (0.33) (Fig. 1 F). Similarly, GFP-SNX4 WT precipitated the HA-SNX32 R220E (1.02), but the amount of precipitant of HA-SNX32 A226E (0.65), HA-SNX32 Q259R (0.75), HA-SNX32 E256R (0.66) and HA-SNX32 R366E (0.49) was comparably less (Fig. 1 F).

### DETERMINANTS OF SUBCELLULAR LOCALIZATION OF SNX32

Previous studies have shown that SNX32 interacts with other SNX-BAR^15^ proteins, but its role in intracellular trafficking was yet to be delineated. To this end, we first quantified the relative amount of SNX32 transcripts by quantitative PCR in different cell lines such as HeLa, U87MG and Neuro2a cells. As shown in Fig S2 A, the expression of SNX32 in HeLa was comparable to brain-derived cells (U87MG, Neuro2a); based on that, we generated a HeLa cell line stably expressing GFP-SNX32 from an inducible promoter of pLVX TRE3G. Upon treatment with a concentration of 500ng/ml of tetracycline for 13hrs, the subcellular localization of SNX32 was punctate. Further, the cells were fixed, immunostained for early endosomal marker (EEA1), Trans Golgi marker (TGN46) and mCherry-Rab11(juxtanuclear recycling endosomal marker), and images were acquired by a laser scanning confocal microscope. As evident from the object-based colocalization calculated using Motion tracking, GFP-SNX32 showed colocalization with early endosomal marker EEA1(Fig.2 A, B), which is also in accord with the study reported by Simonetti. *et al*. ^10^. In addition, we observed a notable amount of SNX32 localized to the cell surface (Fig. 2 C). Also, it showed colocalization with mCherry-Rab11(Fig.2 D) and TGN46 (Fig. 2 E). The partial overlaps were reconfirmed in live-cell microscopy experiments (Supp. video 1) as well as in super-resolution imaging (Fig.2 F-G, Supp. video 2,3).

**Figure 2.**
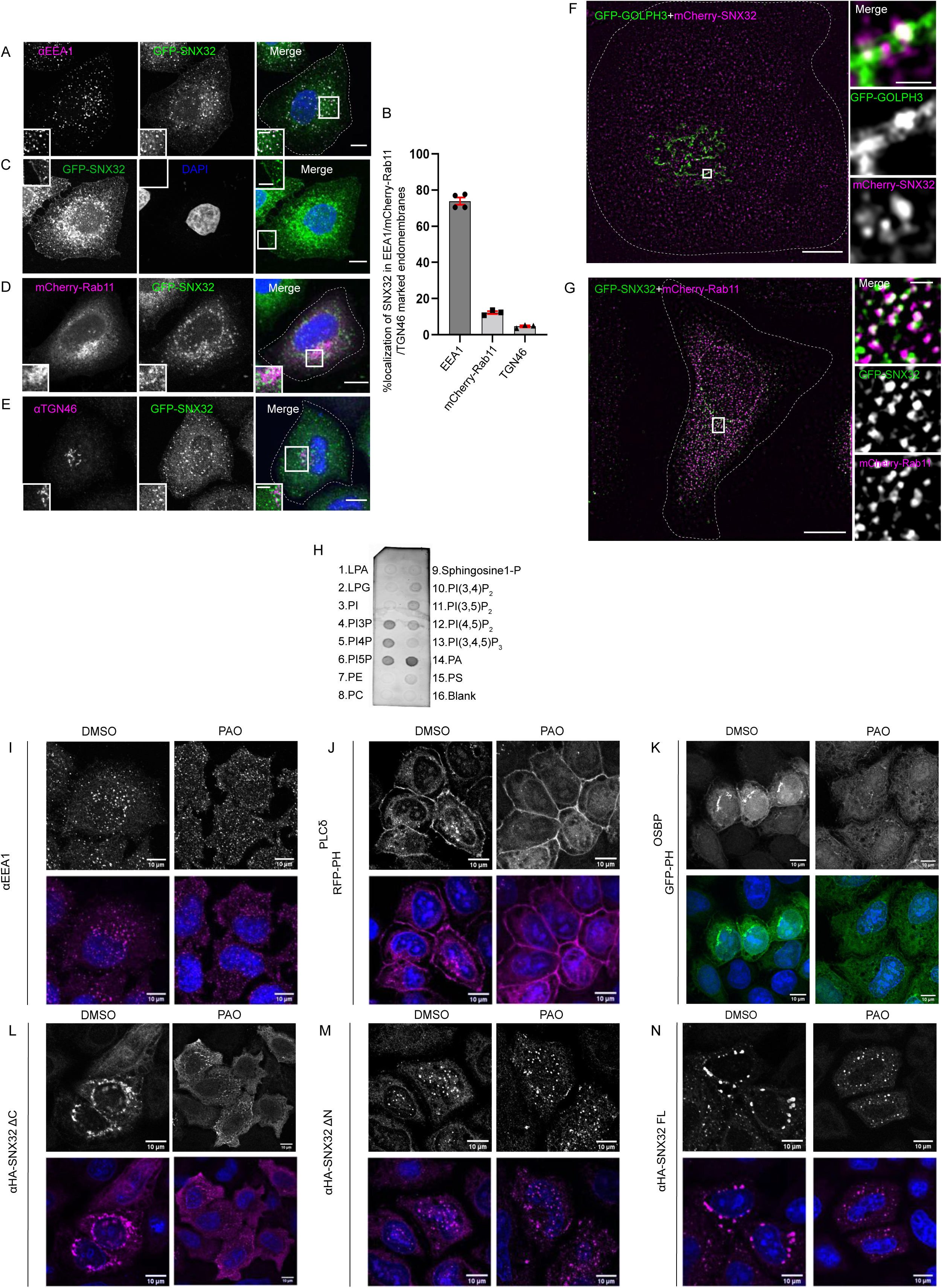
SNX32 undergoes PX domain assisted localization to PI(4)P enriched endosomal membranes in addition to early endosomes. (A) GFP-SNX32 localization with early endosomal marker EEA1, (B) Quantifications showing percentage co-localization of GFP-SNX32 in HeLa cells, data represent mean ±SEM (N=3, n≥60cells per independent experiments) (C) GFP-SNX32 localization at cell periphery (D) GFP-SNX32 colocalization with peri-nuclear recycling endosomal marker mCherry-Rab11, (E) colocalization of GFP-SNX32 with Trans Golgi Network marker TGN46, Scale bar 10µm, inset 5µm (magnified regions are shown as insets). F-G) SIM image showing co-localization of F) GFP-GOLPH3 and mCherry-SNX32, G) GFP-SNX32 and mCherry-Rab11, Scale bar 10µm, inset 1µm (magnified regions are shown as insets). H) PIP Strip membrane immunoblotted using His antibody showing preferential binding of His-SNX32ΔC to PI(3)P, PI(4)P, PI(5)P, PA (representative immunoblot out of 3 biological replicates). I-N) PAO/DMSO treatment in HeLa cells overexpressing I) GFP-PH^OSBP^, J) RFP-PH^PLCį^, K) endogenous EEA1, L) HA-SNX32ΔC, M) HA-SNX32ΔN, N) HA-SNX32FL, Scale bar 10µm.

PX domain was long established as the Phospho inositols (PIns) binding domain, which assists the proteins in preferential membrane association^16^. PIns are short-lived phosphorylated myoinositol heads containing membrane markers showing a stable distribution under normal physiological conditions^21^. Hence, we hypothesized that the PIns affinity of the SNX32 PX domain helps it associate with the endomembrane. Confocal images revealed that HA-SNX32ΔN showed punctate localication scattered through the cytoplasm, and Motion tracking-based analysis did not show colocalization with known endosomal markers (FigS2 B). However, when co-expressed with the interacting partners, GFP-SNX4/GFP-SNX1, they colocalized on EEA1 positive compartments, as quantified using Motion tracking (Fig.S2 C-D). Distinctively, HA-SNX32ΔC was seen as a perinuclear cluster that showed colocalization with PI (4)P binding GFP-PH ^OSBP^ (Fig.S2 E, F), which prompted us to understand the lipid affinity of each domain.

The protein-lipid overlay^22^ assay enables quantitative understanding of the lipid interacting preference of lipid biding proteins. We employed PIP Strips Membranes embedded with 8 phosphoinositide and 7 other biologically relevant lipids. The membrane was incubated with His tagged SNX32ΔC overnight at 4°C. After adequate washes, the membrane was probed for bound His-SNX32ΔC by immunoblotting using an anti-His antibody. The immunoblot of the PIP strip showed that the His-SNX32ΔC was bound to PI(3)P, PI(5)P, PI(4)P, and PA (Fig. 2H). Since the full-length, as well as the SNX32ΔC, was insoluble when expressed in bacteria, we went ahead to validate our observation in HeLa cells using the known inhibitors of lipid kinases. The phosphatidylinositol (PI) kinases-inhibitor assay was carried out utilizing Phenylarsine Oxide (PAO)^23^, a PI4kinase inhibitor, or Wortmannin, a PI3kinase inhibitor^24^, to acutely deplete the intracellular PI(4)P or PI(3)P, respectively^25^. To confirm the specificity of PAO treatment its effect in GFP-PH^OSBP^, a family of tethering proteins that binds specifically to PI(4)P^26^ was assessed. Similarly for confirming the specificity of wortmannin treatment, its effect in EEA1, a FYVE domain containing protein that specifically binds to PI(3)P^27^ was examined. In either inhibitor treatment to negate any of target effect RFP-PH^PLCδ^, a phosphatidylinositol 4,5-bisphosphate (PI(4,5)P2) binding protein^28^ motif was used as a negative control. The cells un-transfected or overexpressing GFP-PH^OSBP^, RFP-PH^PLCδ^, GFP-SNX32, HA-SNX32ΔC, HA-SNX32ΔN were treated with PAO (15ȝM)/ Wortmannin (200nM) in uptake medium for 15mins at 37°C, proceeded for immunofluorescence. The confocal images showed that consistent with the previous reports^25^, PAO treatment caused redistribution of GFP-PH^OSBP^ (Fig.2 I), whereas Wortmannin treatment caused the redistribution of endogenous EEA1(Fig. S2 G) with less or no effect on the distribution of other PIns binding markers^25^, indicating the depletions were specific on either case (Fig.2 J-K, Fig.S2 H-I). In line with the Lipid overlay assay and localization studies, the PAO treatment caused redistribution of HA-SNX32ΔC from the perinuclear cluster to the cytosol (Fig.2 L) with less or no effect on the localization of HA-SNX32ΔN (Fig.2 M). In contrast, wortmannin treatment caused HA-SNX32ΔN to relocalize from the punctae to the cytosol (Fig. S2 J) with less effect on HA-SNX32ΔC (Fig. S2 K). For both the inhibitor treatment, the altered localization of the SNX was less pronounced in the case of the full-length protein (Fig.2 N, Fig.S2 L). The effect of PAO on HA-SNX32ΔC was confirmed using a membrane fractionation experiment^29^. HeLa cells overexpressing GFP-PH^OSBP^ or HA-SNX32ΔC was incubated with PAO for 15mins at 37°C followed by snap-freezing to fracture the membrane and release the cellular contents. The membrane and cytosolic fractions(S) were separated by centrifugation, further, the membrane fraction was solubilized with detergent to obtain a pellet fraction(P). The fractions (S, P) were resolved through SDS-PAGE and subjected to immunoblotting using an anti-GFP, anti-HA,anti-Vinculin, anti-TfR antibody, where TfR is a membrane protein indicating the purity of the fractions. We found that the distribution of HA-SNX32ΔC was increased in the cytosolic fraction when treated with PAO in comparison to the DMSO treated condition (Fig.S2 M).

### SNX32 REGULATES TRANSFERRIN TRAFFICKING

In order to understand the functional significance of SNX32 and SNX4 association, we sought to see if the absence of SNX32 in the system shows any effect on cargo trafficking. Previously, it has been reported that SNX4 suppression alters the levels of Transferrin Receptor, TfR ^30^, by diverting the cargo molecules from the recycling route to the lysosomal degradative pathway. Though SNX32 is largely colocalised with SNX4 on EEA1-positive early endosomal compartments (Fig. 3 A), we also observed a small but significant SNX32 population on the transferrin receptor-positive recycling compartment (Fig.3 B). Our results from confocal microscopy on paraformaldehyde-fixed cells and live-cell video microscopy revealed that SNX32 and SNX4 colocalize (Fig. 3 C, D) and co-transports with Transferrin (Supp. video 4). We, therefore, assessed the recycling of Transferrin in HeLa cells depleted for SNX32(SNX32KD) using SMART pool siRNA (Fig S3 A) or shRNA clones (shSNX32, Fig S3 B). HeLa cells treated with SCR, SNX32KD were serum starved for 2hrs, incubated with 10µg/ml of transferrin (Alexa Fluor 488 conjugated) for 30min Pulse and chased using 100µg/ml of holotransferrin for a maximum of 60mins at 37°C. It was observed that in SNX32 silenced condition, compared to Scramble, there was no significant difference in integral intensity of transferrin in the initial time points(0-10min) of the chase period. Though in the course of a 60min chase, the scrambled cells followed a trend of diminishing integral fluorescence intensity (Fig.3 E), whereas SNX32 suppressed cells retained the intensity (Fig.3 E). We characterized the nature of the endosomes in which transferrin was present and observed that it was retained in the early endosome positive for EEA1(Fig.3 F, G). Moreover, the receptor’s distribution along with the transferrin (Alexa Fluor 568 conjugated) revealed both being present in the EEA1 positive vesicles (Fig.3 H, I). In agreement with this, the shRNA-mediated SNX32 downregulated conditions (Fig. 3 J, K) also showed similar effects. We further validated the effect of SNX32 downregulation in the early endosomal redistribution of transferrin by compensating for the loss of SNX32. We observed that by overexpressing the shRNA resistant SNX32(HA-shSNX32#4r), in the background of shSNX32#4 mediated SNX32KD, the colocalization of transferrin (Alexa Fluor 568 conjugated) with EEA1 was reduced; however, the overexpression of the individual domains (HA-SNX32ΔC, HA-SNX32ΔN#4r) failed to show any such effect, suggesting their role in cargo trafficking (Fig S3 C, D). It was previously reported that Cation Independent Mannose-6-Phosphate Receptor (CIMPR) interacts with SNX32^1^ and the interaction follows the canonical sequence-dependent recognition similar to CIMPR and SNX5/SNX6^10^. Even though CIMPR is shown to be a potential cargo for SNX32^1, 10^ the involvement of SNX32 in the trafficking of CIMPR was not yet reported; additionally, it has been shown that the suppression of SNX1 leads to the degradation of CIMPR^31^. Our results from confocal microscopy showed that SNX32 and SNX1 colocalize (Fig.S3 E, F) and co-transports CIMPR (Supp. video 4). Further, we observed that the SNX32KD resulted in the steady-state redistribution of CIMPR (Fig.S3 G, H), which was also seen to be correspondingly more with EEA1 positive early endosomes (Fig.S3 G, H). We did not find any significant reduction in the total level of CIMPR upon suppression of SNX6/SNX32 (Fig.S3 I, J). To further confirm whether the steady-state redistribution of CIMPR upon SNX32KD resulted from a defect in the kinetics of transport, we performed antibody uptake assay utilizing CD8-CIMPR reporter endocytosis^32^. In control cells, after incubating the cells with an anti-CD8 antibody to label the cell surface CD8-CIMPR, the kinetics of transport revealed that the CD8-CI-MPR had reached Golgi after an incubation period of 30 minutes (Fig.S3 K, L). In contrast, the HeLa cells suppressed for SNX32, the internalized anti-CD8 antibody was observed to be accumulated in endosomes and were unable to reach Golgi after 30 minutes of chase period (Fig.S 3 K, L)

**Figure 3.**
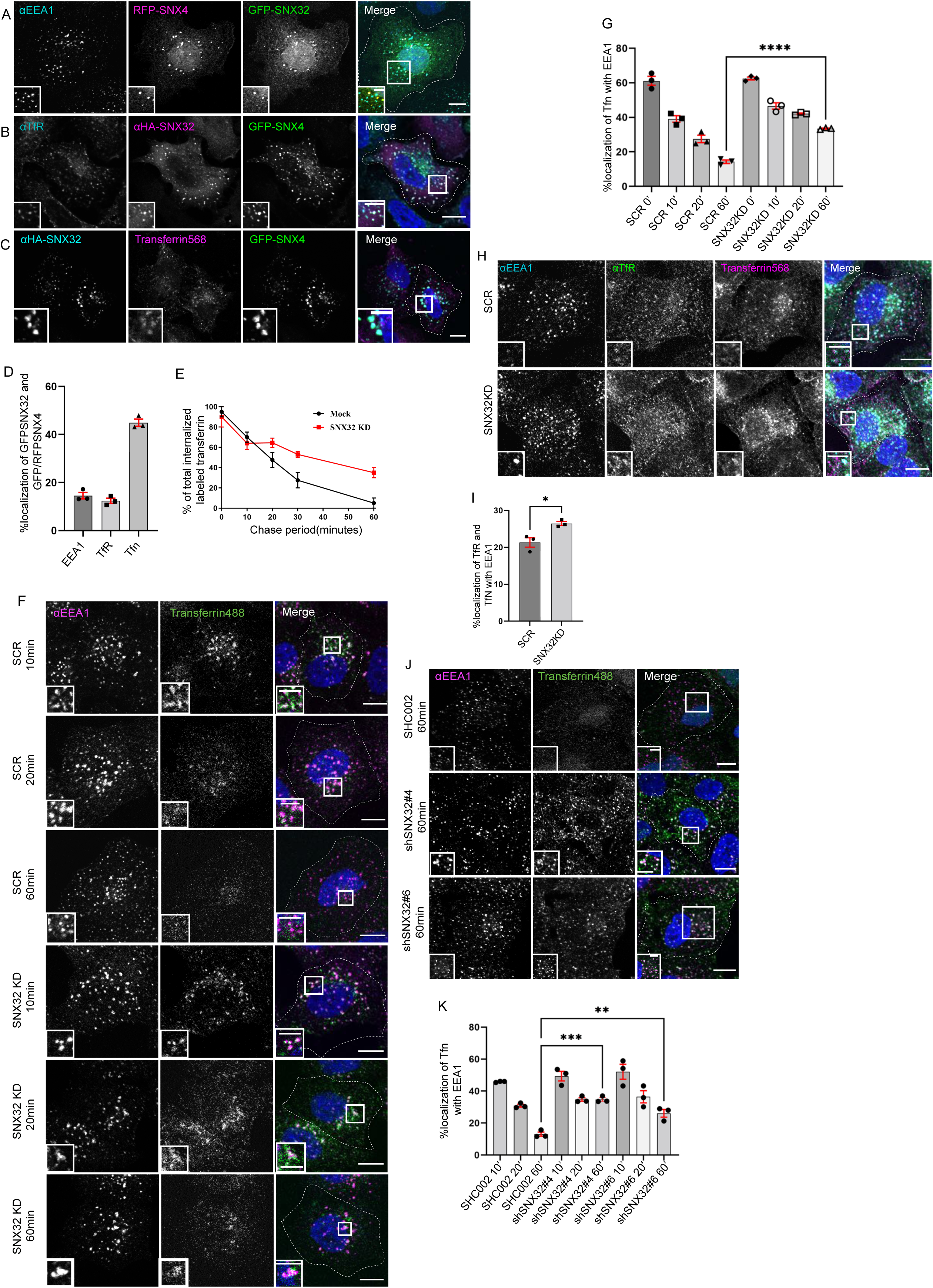
SNX32 interacts with and regulate the trafficking of transferrin bound transferrin receptor in HeLa cells. A-D) SNX32 and SNX4 colocalises with (A) early endosome marker EEA1, (B) peri-nuclear recycling endosomal marker TfR, (C)Alexa Fluor 568 conjugated transferrin, Scale bar 10µm, inset 5µm (magnified regions are shown as insets), D) Percentage localization of SNX32 and SNX4 with EEA1, TfR and Transferrin, data represent mean “SEM (N=3, n≥60cells per independent experiments). E) Quantitative estimation of the percentage of total internalized transferrin (Alexa Fluor 488 conjugated) in different chase periods measured by its total intensity of fluorescence in mock and SNX32 suppressed cells, data represent mean ±SEM (N=3, n≥60cells per independent e[periments). F) Followed by SMARTpool mediated gene down regulation of Scrambled (SCR) or SNX32, transferrin (Alexa Fluor 488 conjugated) Pulse-Chase experiment was carried out as described in materials and method section, the cells were fixed at specified timepoints, immunostained using early endosomal marker EEA1, DAPI was used to stain nucleus, Scale 10µm, inset 5µm (magnified regions are shown as insets). G) Quantification of percentage localization of transferrin (Alexa Fluor 488 conjugated) with EEA1 at corresponding time points, data represent mean “SEM (N=3, n≥60cells per independent experiments), P value <0.0001 Unpaired t test Two-tailed (**** P < 0.0001). H)Followed by SMARTpool mediated gene down regulation of Scrambled (SCR) or SNX32, HeLa cells were allowed to uptake Transferrin (Alexa Fluor 568 conjugated) for 30min, 37°C, fixed and immunostained using EEA1, TfR, DAPI was used to stain nucleus, Scale 10µm, inset 5µm (magnified regions are shown as insets).I) Quantification of percentage localization of TfR with internalised transferrin and EEA1, data represent mean “SEM (N=3, n≥60cells per independent experiments), P value 0.0206 (* P < 0.05), Unpaired t test Two-tailed. J)Followed by shRNA mediated gene down regulation of Scrambled (SHC002) or SNX32 shRNA clones (shSNX32#4,shSNX32#6) transferrin (Alexa Fluor 488 conjugated) Pulse-Chase experiment was carried out, the cells were fixed at timepoints specified, immunostained using EEA1, DAPI was used to stain nucleus, Scale 10µm, inset 5µm(magnified regions are shown as insets), K) Quantification of percentage localization of internalised transferrin with EEA1 at corresponding timepoints, data represent mean “SEM (N=3, n≥60cells per independent experiments), P value 0.0003(** P < 0.01, *** P < 0.001), One way ANOVA, Dunnett’s multiple comparisons test.

Since SNX6 is the closest homolog of SNX32, we investigated if SNX6 downregulation (SNX6KD) also causes effects in transferrin trafficking. As previously described, the transferrin Pulse-Chase was carried out in SCR/SNX6KD cells. Comparable to SNX32KD, SNX6KD also displayed that the transferrin (Alexa Fluor 488 conjugated) was retained at EEA1 positive early endosome even after 30min of chase period (Fig.S4 A, B). We then asked if SNX32 and SNX6 have any overlapping roles in transferrin trafficking. To address this, we overexpressed GFP-SNX32 in SNX6KD cells and measured the transferrin (Alexa Fluor 568 conjugated) colocalizing with the EEA1 positive compartment. As observed in Fig S4 C-D, the colocalization of transferrin with EEA1 is significantly reduced in the SNX6KD cells expressing GFP-SNX32. We also observed that the downregulation of SNX32 or SNX6 did not alter the total TfR levels in HeLa cells (Fig.S3 E, F).

### INTERPLAY OF SNX32 WITH SNX4 AND RAB11 IN TRANSFERRIN TRAFFICKING

Next, to have a better understanding of the SNX4 and SNX32 regulated transferrin trafficking pathways, we investigated transferrin trafficking in double knockdown conditions. HeLa cells treated with SCR, siSNX4 and/or siSNX32 were serum starved for 2hrs, incubated with 10µg/ml of transferrin (Alexa Fluor 568 conjugated) for 30min Pulse and chased using 100µg/ml of holotransferrin for a maximum of 30mins at 37°C. As reported earlier ^30^, the downregulation of SNX4(SNX4KD) resulted in increased colocalization of transferrin in EEA1 positive compartment(Fig. 4 A); moreover, the double downregulation of SNX4 and SNX32 didn’t show any significant difference in the percentage of transferrin colocalised with EEA1 marked early endosome(Fig. 4 B). Further, we went ahead to examine if SNX32 can compensate for the loss of SNX4. It was observed that in SNX4KD cells overexpressing GFP-SNX32 did not show any significant change in the transferrin colocalization with the EEA1 compartment (Fig. 4 C, D). Based on the above observations, we propose that SNX4 and SNX32 may function in a common pathway regulating the intracellular trafficking of transferrin. The physical association of SNX4 and SNX32 may provide an explanation for the above observations.

**Figure 4.**
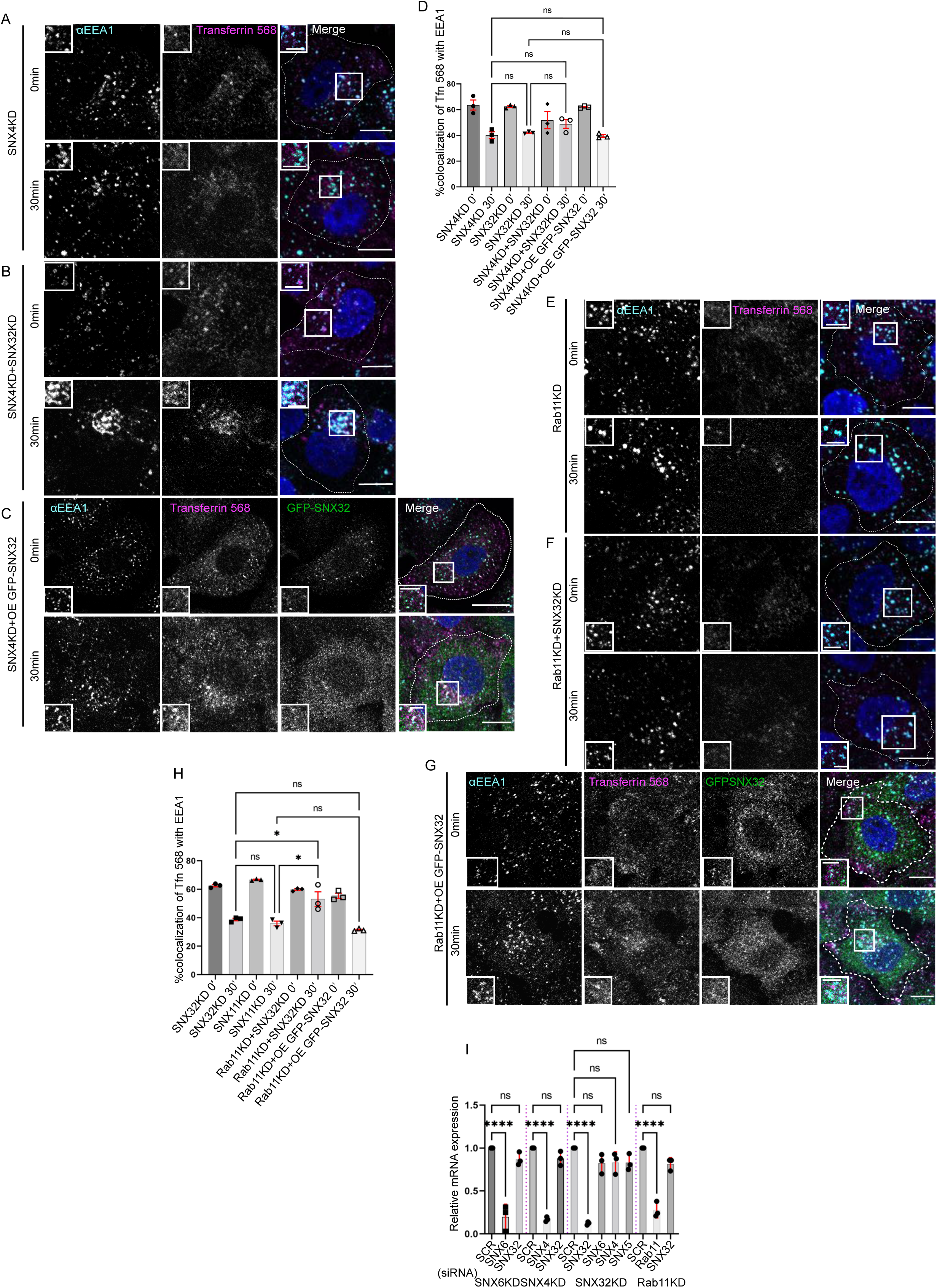
Interplay of SNX32, SNX4 and Rab11 in transferrin trafficking: A-C) Followed by SMARTpool mediated gene down regulation of (A)SNX4, (B)SNX4 and SNX32 (C) SNX4KD and over expression of GFP-SNX32, the transferrin (Alexa Fluor 568 conjugated) Pulse-Chase experiment was carried out as described in materials and method section, the cells were fixed at specified timepoints, immunostained using early endosomal marker EEA1, DAPI was used to stain nucleus, Scale 10µm, inset 5µm (magnified regions are shown as insets).D) Quantification of percentage localization of transferrin (Alexa Fluor 568 conjugated) with EEA1 at corresponding time points, data represent mean ±SEM (N=3, Q≥15 random frames per independent experiments), P value <0.0636, Ordinary one-way ANOVA, âtdik’V PXOWiSOe comparisons test (ns-nonsignificant). E-G) Followed by SMARTpool mediated gene down regulation of (E) Rab11, (F) Rab11 and SNX32 (G) Rab11KD and over expression of GFP-SNX32, transferrin (Alexa Fluor 568 conjugated) Pulse-Chase experiment was carried out as described in materials and method section, the cells were fixed at specified timepoints, immunostained using early endosomal marker EEA1, DAPI was used to stain nucleus, Scale 10µm, inset 5µm (magnified regions are shown as insets). H) Quantification of percentage localization of transferrin (Alexa Fluor 568 conjugated)) with EEA1 at corresponding time points, data represent mean ±SEM (N=3, Q≥15 random frames per independent experiments), P value <0.0030, Ordinary one-way ANOVA, Šídák’s multiple comparisons test (* P < 0.05, ns-nonsignificant). I) Analysis of the relative gene expression levels of SNX5, SNX6, SNX4 and SNX32 by qRT-PCR in HeLa cells depleted for SNX4, SNX6, SNX32 and Rab11. Values of control were arbitrarily set as 1 against which experimental data were normalized. Gapdh was used as internal control, data represent mean ±SEM (N=3) P value <0.0001, Ordinary one-way ANOVA, Šídák’s multiple comparisons test (**** P < 0.0001, ns-nonsignificant).

Since Rab11 is an established regulator of transferrin trafficking, we next assessed weather the GTPase plays any role in SNX32-regulated TfR trafficking. HeLa cells treated with SCR, siRab11 and/or siSNX32 cells were serum starved for 2hrs, incubated with 10µg/ml of transferrin (Alexa Fluor 568 conjugated) for 30min Pulse and chased using 100µg/ml of holotransferrin for a maximum of 30mins at 37°C. As reported earlier, the colocalization of transferrin with EEA1 was significantly higher in Rab11 downregulated (Rab11KD) cells compared to the control cells (Fig. 4 E). Further, the depletion of both proteins led to a significant increase in the localization of transferrin with EEA1 compared to either SNX32KD or Rab11KD cells (Fig. 4 F). Moreover, it was observed that the overexpression of GFP-SNX32 in Rab11KD cells did not show any significant difference in the transferrin colocalization with EEA1(Fig. 4 G, H). The above observations are suggestive of the existence of an independent pathway, in addition to the one in which SNX32 may play a role upstream of Rab11.

We also noted that the downregulation of SNX32 did not have any effect on the mRNA level of either of SNX4, SNX5, SNX6 or Rab11. Similarly, the knockdown of SNX6, SNX4 or Rab11 did not alter the mRNA expression level of SNX32(Fig. 4 I).

### SNX32 INTERACTS WITH THE CARGO THROUGH ITS PX DOMAIN

Next, we sought to understand the selectivity of association of the individual interacting partner, SNX32 and SNX4/SNX32 and SNX1, with the cargo TfR/CIMPR. We performed a GBP-based immunoprecipitation assay where GST-GBP was immobilized on GST beads and incubated with HeLa cell lysate overexpressing GFP-SNX4 WT/GFP-SNX1 WT/GFP-SNX32 WT. The eluates were resolved through SDS-PAGE and subjected to immunoblotting using an anti-GFP and anti-TfR or anti-CIMPR antibody. We observed that GFP-SNX32 was efficiently precipitating TfR (1.2) as well as CIMPR (0.38), however, TfR precipitated with GFP-SNX4 (0.59) and CIMPR precipitated with GFP-SNX1 (0.12) was comparably less (Fig. 5 A, B). Earlier reports showed that SNX5 interacts with its cargo, CIMPR^1^/IncE ^33^, through its PX domain. To explore weather SNX32 also interacts with the cargo through its PX domain, we employed the His-tagged deletion mutant of SNX32, His-SNX32ΔC(1-166aa) (Fig. S1 A). Affinity chromatography-based pulldown assay was carried out by immobilizing His-SNX32ΔC on Ni-NTA beads and incubating with an enriched membrane fraction of HeLa cell lysate. The eluates were resolved through SDS-PAGE and subjected to immunoblotting using an anti-His and anti-TfR or anti-CIMPR antibody. Using the affinity chromatography-based pulldown assay, we showed that the PX domain of SNX32 contributes to the interaction with TfR/CIMPR (Fig. 5 C, D).

**Figure 5.**
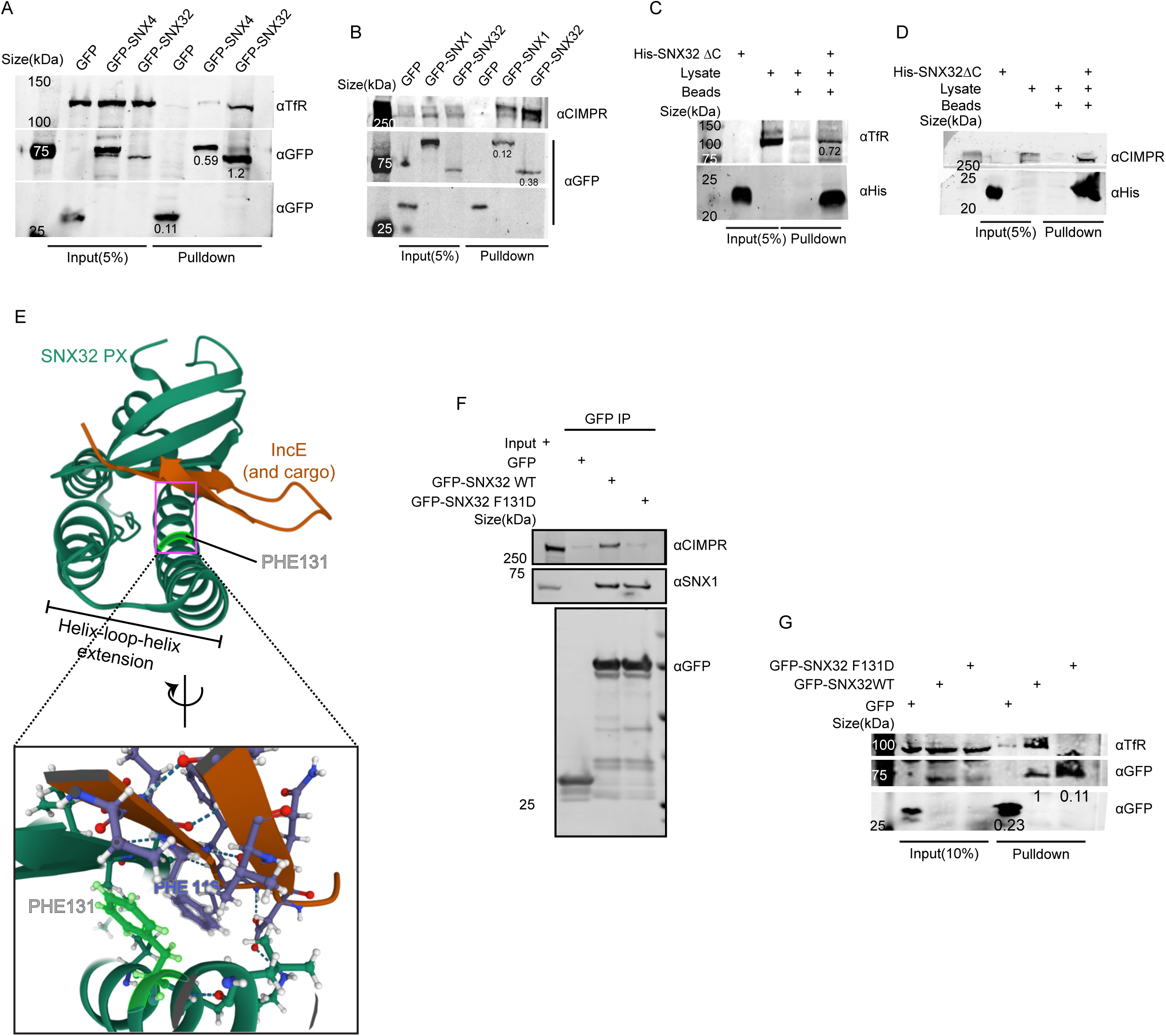
PX domain of SNX32 participates in the interaction with CIMPR, TfR and F131 of SNX32 is critical for interaction with cargo: A) GBP co-immunoprecipitation of GFP tagged SNX4 and SNX32 transiently transfected in HEK293T cells showing GFP-SNX32 efficiently precipitating TfR, GBP immunoprecipitation was carried out as described in materials and methods section and immunoblotted using GFP and TfR antibody (representative immunoblot out of 3 biological replicates, values represent the ratio of TfR to GFP band intensity). B) GBP co-immunoprecipitation of GFP tagged SNX1 and SNX32 transiently transfected in HEK293T cells showing GFP-SNX32 efficiently precipitating CIMPR, GBP immunoprecipitation was carried out as described in materials and methods section and immunoblotted using GFP and TfR antibody (representative immunoblot out of 3 biological replicates, values represent the ratio of CIMPR to GFP band intensity).C) His affinity chromatography-based pulldown showing His-SNX32ΔC precipitating TfR from membrane enriched HeLa cell lysate fraction, His pulldown was carried out as described in material method section and immunoblotted using His and TfR antibody (representative immunoblot out of 3 biological replicates, values represent the ratio of His to TfR band intensity). D) His affinity chromatography-based pulldown showing His-SNX32ΔC precipitating CIMPR from membrane enriched HeLa cell lysate fraction, His pulldown was carried out as described in material method section and immunoblotted using His and TfR antibody (representative immunoblot out of 3 biological replicates). E)Cargo/IncE binding site of SNX32 PX domain as observed in crystal structure (PDB ID:6E8R) reported by Chandra et.,al^16^, inset showing the stacking interaction between F131 of SNX32 and F116 of IncE F) Co-immunoprecipitation of GFP tagged SNX32 wild type (WT) and GFP trap of GFP-tagged SNX32 WT/ SNX32 F131D, showing both efficiently precipitating ESCPE-1 sub unit SNX1 whereas SNX32 F131D failed to precipitate CIMPR, each transiently transfected in HEK293T cells. The elute was resolved in SDS-PAGE and immunoblotted using GFP, SNX1 and CIMPR antibody G) GBP co-immunoprecipitation of GFP tagged SNX–proteins transiently transfected in HeLa cells showing GFP-SNX32 but not GFP-SNX32 F131D efficiently pulling down TfR, GBP immunoprecipitation was carried out as described in materials and methods section and immunoblotted using GFP and TfR antibody (representative immunoblot out of 3 biological replicates, values represent the ratio of TfR to GFP band intensity).

As previously reported, the PX domains of SNX32, SNX6 and SNX5 contain a conserved stretch of 38 amino acids^34^ (Fig. S5 A) that, with a helix-loop-helix fold, provides a hydrophobic groove for binding to the bipartite sorting motif present in cargo proteins that include CIMPR^1^. Moreover, the conserved phenylalanine within the stretch was found crucial for maintaining the interaction between SNX5 and CIMPR^1^ without influencing the dimerization with its heterodimerization partner, SNX1/SNX2^1^. We superimposed the already available crystal structure of IncE bound SNX32 PX domain^16^ on CIMPR bound SNX5^1^ structure. The analysis of the superimposed structure revealed that the conserved phenylalanine (F131 for SNX32) engages in a stacking interaction with a Phe residue of IncE (Fig. 5 E). Of note, the Inc proteins of the pathogen are known to mimic the interactions of native host proteins with the sorting machinery to establish a successful infection^33^. Thus, the F131 of SNX32 (Fig. S5 B) is predicted to directly associate with cargo proteins containing the ESCPE-1 bipartite sorting motif. To delineate the importance of the residue, we performed a GBP-based immunoprecipitation assay, where GST-GBP was immobilized on GST beads and incubated with HeLa cell lysate overexpressing GFP-SNX32 WT/GFP-SNX32 F131D. The eluates were resolved through SDS-PAGE and subjected to immunoblotting using an anti-GFP and anti-CIMPR antibody. As shown in Fig. 5 F, the SNX32 F131D mutant failed to precipitate CIMPR, whereas it was successful in precipitating endogenous SNX1, thereby validating the role of the Phe in binding with CIMPR. We further investigated if the interaction between SNX32 and TfR also follows a similar mechanism. The GST-GBP immunoprecipitation of GFP-SNX32 WT/GFP-SNX32 F131D showed that the F131D mutant of SNX32 failed to precipitate TfR (Fig 5 G). The observation affirmed the role of F131 in TfR binding.

### SNX32 PLAYS A CRUCIAL ROLE IN REGULATING NEURITE OUTGROWTH

To understand the functional significance of SNX32, we focused on neuronal differentiation. Neurite outgrowth formation, projection, and extension of neural processes are fundamental in normal neuronal development. During neuronal differentiation and polarity establishment, the directional intracellular transport maintains the supply and distribution of molecules to the growing processes^35^. Neurite outgrowth assay^36^ serves as an *in vitro* model to investigate the potential effects of any particular molecule of interest in neuronal differentiation.

We used ON-TARGETplus SMARTpools for targeted siRNA-mediated downregulation of SNX32 (SNX32KD), and SNX6 (SNX6KD), with siRNA-scrambled (SCR) as a negative control (Fig. S5 C). SCR/SNX32KD/SNX6KD Neuro-2a cells were differentiated to form neurites by replacing the standard growth media (MEM with 10%FBS) with the differentiation media (MEM with 1%FBS containing 10 µmol/L Retinoic acids), and incubation was continued for the next 48hrs. Further, the cells were fixed, and phase contrast images were acquired by AxioVert.A1 microscope. Neurite length was measured using ImageJ software, and only neurites measuring more than the diameter of the cell body in length were taken for final analysis. We observed that in SNX32 depleted (SNX32KD) Neuro2a cells, a mouse neuroblastoma cell line, the number of cells with elongated neurites was less compared to SCR or SNX6KD conditions (Fig. 6 A, B). Even though most of the cells in the SNX32KD condition showed positive sprouting, the length of these sprouts was limited (less than the diameter of the cell body). To better understand and negate any artefacts introduced due to cell fixation, the procedure of neurite induction was replicated, and images were captured in real time for a maximum of 48hrs. The DIC time-lapse live-cell video imaging (Supp. video 5) showed that response to retinoic acid (RA) supplemented in 1% FBS containing MEM media was prompt and rapid. Sprouting and normal growth cone movement were visible in ≥ 80% of the cells within the initial few hours in all the conditions under investigation (Fig. 6 C-E). As time progressed, in the case of SCR and SNX6KD long neurites became more prevalent and intermingled with adjacent cell neurites, forming intricate networks (Fig. 6 F-G). Whereas in the case of the SNX32KDcondition, the sprouts failed to elongate and were inefficacious in establishing the neural network (Fig. 6H). However, it is interesting to note that in either case of SNX32KD/SNX6KD conditions, cell proliferation was much slower than in the SCR condition (Supple. video1). To have a better insight into the role of SNX32 in neurite outgrowth, we applied a SILAC-based proteomics approach, hypothesizing that the role of SNX32 in neuronal development could be attributed to its cargo repertoire^10^.

**Figure 6.**
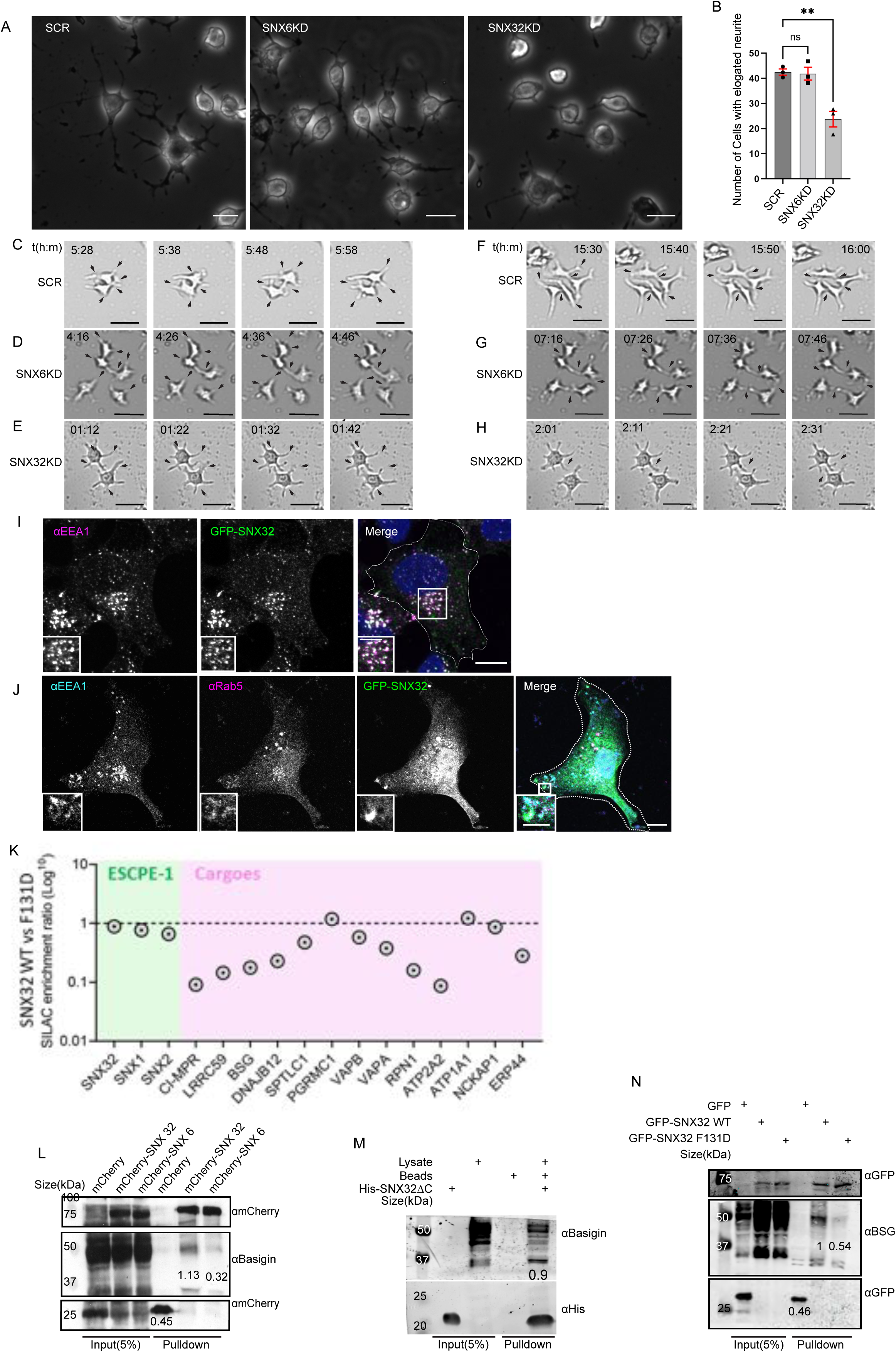
SNX32 plays a crucial role in neurite differentiation, and it interacts with immunoglobulin superfamily member BSG via its PX domain: A) Phase contrast image of Neuro2a cells transfected with Scrambled (SCR)/ SNX32/ SNX6 siRNA SMARTpool followed by neurite induction as described in the materials and methods section, fixed, and imaged using Zeiss Axio vert. A1 microscope, Scale bar 50µm. B) Quantification of number of cells with elongated neurites, data represent mean ±SEM (N=3, n≥100cellV per independenW experiment, values are means ± SEM), P value 0.0025(** P < 0.01), Ordinary one-way ANOVA Dunnett’s multiple comparisons test. C-E) Snap Shots of SCR/SNX6/SNX32 SMARTpool siRNA transfected Neuro2a cells induced with RA for neurite induction showing neurite sprouting, Scale bar 50 µm (black arrows pointing to sprouting). F-H) Snap Shots of Neuro2a cells transfected with SCR/SNX6/SNX32 SMARTpool siRNA transfected Neuro2a cells induced with RA for neurite induction showing neurite extension, black arrows pointing network formation/retraction of neurites, Scale bar 50 µm. I) SHSY5Y cells showing colocalization of GFP tagged SNX32 with EEA1 harbouring endosomes, Scale 10µm, inset 5µm (magnified regions are shown as insets). J) U87MG cells showing the colocalization of GFP tagged SNX32 with Rab5 and EEA1 harbouring endosomes, Scale 10µm, inset 5µm (magnified regions are shown as insets). K) SILAC enrichment ratio (Log10) of SNX32 WT vs SNX32 F131D, SILAC was carried as described in the materials and methods sections, the proteins consistently appeared in at least 2 experimental replicates were considered for the final analysis. L) Co-immunoprecipitation of mCherry tagged SNX–proteins transiently transfected in U87MG cells showing mCherry SNX32 but not mCherry SNX6 efficiently pulling down Basigin (BSG), mCherry nanobody mediated pulldown was carried out as described in materials and methods section and immunoblotted using mCherry and BSG antibody (representative immunoblot out of 3 biological replicates, values represent the ratio of BSG to mCherry band intensity). M) Histidine (His) pulldown showing His SNX32ΔC efficiently pulling down BSG from membrane enriched HeLa lysate (representative immunoblot out of 3 biological replicates, values represent the ratio of BSG to His band intensity). N) GFP tagged SNX–proteins transiently transfected in HeLa cells showing GFP SNX32 WT but not GFP SNX32 F131D efficiently pulling down Basigin (BSG), GFP nanobody mediated pulldown was carried out as described in materials and methods section and immunoblotted using GFP and BSG antibody (representative immunoblot out of 3 biological replicates, values represent the ratio of BSG to GFP band intensity).

### DIFFERENTIAL PROTEOMICS IMPLIED BSG AS AN INTERACTING PARTNER OF SNX32

To provide mechanistic insight into the role of SNX32, we performed quantitative Stable isotope labelling by amino acids in cell culture (SILAC)-based proteomics on defining the SNX32 interactome. To establish this methodology, we first lentivirally transduced SHSY5Y, a human neuroblastoma cell line to stably express GFP-tagged SNX32. In agreement with the previous report in RPE1 cells^10^, in SHSY5Y cells, GFP-SNX32 largely colocalized with EEA1 harbouring endosomes (Fig. 6 I). The endosomal localization of GFP-SNX32 with EEA1 and in addition, Rab5 was also observed in U87MG (Fig.6 J).

The earlier study^10^ in RPE1 cells revealed that several proteins implicated in neuronal differentiation, including Basigin (BSG)^37^, PGRMC1^38^, ROBO1^39^, and SV2a^40^, were enriched in the interactome of SNX32 but not the interactome of SNX6^10^. We next extended this analysis through differential SILAC-based proteomics by comparing the interactomes of wild-type SNX32 and SNX32 F131D. As previously described, the F131D mutant of SNX32 fails to interact with cargoes such as CIMPR (Fig. 5 F) and TfR (Fig. 5 G). We further established an SHSY5Y cell line stably expressing GFP-SNX32 F131D. It was observed that GFP-SNX32 F131D also retained the ability to associate with endosomes (Fig. S5 D).

This comparative proteomics (Fig. S5 E) highlighted several cargoes that bind to SNX32 through a mechanism that requires the critical F131 residues within the hydrophobic groove of the SNX32 PX domain. Based on the Abundance Ratio: (SNX32 F131D) / (SNX32 WT), we found several transmembrane proteins interacting with SNX32, including the well-established cargo CIMPR. Among these, BSG was one of the strongest quantified interactors in addition to CIMPR. BSG showed enrichment in SNX32 interactome but a drastic reduction in F131D interactome (Fig. 6 K). BSG is a transmembrane receptor belonging to the superfamily of immunoglobulins. BSG, also known as EMMPRIN, CD147, or human leukocyte activation-associated M6 antigen, is involved in a myriad of cellular processes, including tissue remodelling, visual sensory system development in mice, regulation of neuronal development and the cell surface concentration of monocarboxylate transporters^41^. We utilized a mCherry nanobody-based immunoprecipitation (mCBP)^42^ assay for the same. The GST-mCBP immunoprecipitation assay was carried out by immobilizing GST-mCBP on GST beads and incubating with U87MG cell extracts overexpressing mCherry/mCherry-SNX6/mCherry-SNX32. The eluates were resolved through SDS-PAGE and subjected to immunoblotting using an anti-mCherry and anti-BSG antibody. Immunoblotting using an anti-BSG antibody showed that endogenous BSG was efficiently precipitated by mCherry-SNX32 but not mCherry-SNX6(Fig.6 L). As previously demonstrated, the PX domain of SNX32 precipitates CIMPR and TfR. To identify if a similar mechanism is relevant in the case of interaction with BSG, we performed an affinity chromatography-based pulldown assay. The assay was carried out by immobilizing His-SNX32ΔC (Fig. S1 A, C) on Ni-NTA beads and incubating with an enriched membrane fraction of HeLa cell lysate. The eluates were resolved through SDS-PAGE and subjected to immunoblotting using an anti-His and anti-BSG antibody. Using the affinity chromatography-based pulldown assay, we showed that the PX domain of SNX32 contributes to the interaction with BSG (Fig. 6 M). We also carried out GFP-nanobody-based coimmunoprecipitation of full-length GFP-SNX32 WT and GFP-SNX32 F131D. While the endogenous BSG protein could be coimmunoprecipitated with the GFP-SNX32 wild type, the interaction was lost for the mutant (Fig.6 N). Together, these data establish BSG as a selective interacting partner for canonical-binding to the PX domain of the ESCPE-1 component SNX32.

### SNX32 PHENOCOPIES BSG IN NEURITE OUTGROWTH ASSAY

Previously, it was reported that BSG plays a crucial role in complex^43^ and space-filling dendrite growth^44^ in *Drosophila*. As we observed a similar phenotype in SNX32 suppressed conditions, we next set forth to see if the case is similar under BSG downregulation in neurite induction of Neuro2a cells. Here, we used shRNA clones of the MISSION shRNA product line from Sigma Aldrich to downregulate SNX32(SNX32KD) or BSG(BSGKD) in Neuro2a cells. The TRC1.5 pLKO.1-puro non-Mammalian shRNA Control Plasmid DNA (SHC002) was used as a negative control containing a sequence that should not target any known mammalian genes but still engage the RISC complex. The knockdown efficiency was measured at the mRNA level using qRT-PCR (Fig. S6 A-B). The SNX32KD/BSKD cells were differentiated to form neurites by replacing the standard growth media (MEM with 10%FBS) with the differentiation media (MEM with 1%FBS containing 10 µmol/L Retinoic acid) and incubated for the next 48hrs. Further, the cells were fixed, and phase contrast images were acquired by AxioVert.A1 microscope. Neurite length was measured using ImageJ software, and only neurites measuring more than the diameter of the cell body in length were taken for final analysis. It was observed that in BSG deficient cells, the number of cells with elongated neurites was less, which is in accordance with the previous reports^37^. Further, the number of cells with elongated neurites in the case of SNX32KD was also comparable to that of BSGKD (Fig. 7 A, B, Supp. video 6). The observation suggests that the gene silencing of SNX32 phenocopied that observed in BSG deficient cells^37^. Most cells showed sprouting in either SNX32 or BSG-suppressed conditions, but the length of these sprouts was limited (less than the diameter of the cell body) compared to the non-targeting condition (SHC002). To better understand and negate artefacts due to cell fixation, the procedure of neurite induction was replicated, and images were captured in real-time for a maximum of 48hrs (Supp. video 6). The DIC time-lapse live-cell video imaging showed that the response to RA supplemented in 1% FBS containing MEM media was prompt and rapid. Sprouting was visible in • 80% of the cells within the initial few hours in all conditions (Fig. 7 C-G). In control cells, long neurites were prevalent, and they intermingled with adjacent cell’s neurites to form intricate networks (Fig. 7 H). In contrast, upon SNX32 or BSG suppression mediated by shRNA clones shSNX32#4 or shBSG#7, sprouts failed to elongate, and they were inefficacious in establishing the neural network (Fig. 7 I, J), while in cells transfected with shSNX32#6 or shBSG#8 the neural tubes elongated but without branching or the formation of any network (Fig. 7 K, L). Interestingly, in neither case, the growth cone movement was affected (Supp. video 6), which is consistent with the perturbation of neurite extension observed upon treatment with an antibody against BSG^37^. Taken together, these observations suggest that SNX32 is functionally linked with BSG in neuronal development.

**Figure 7.**
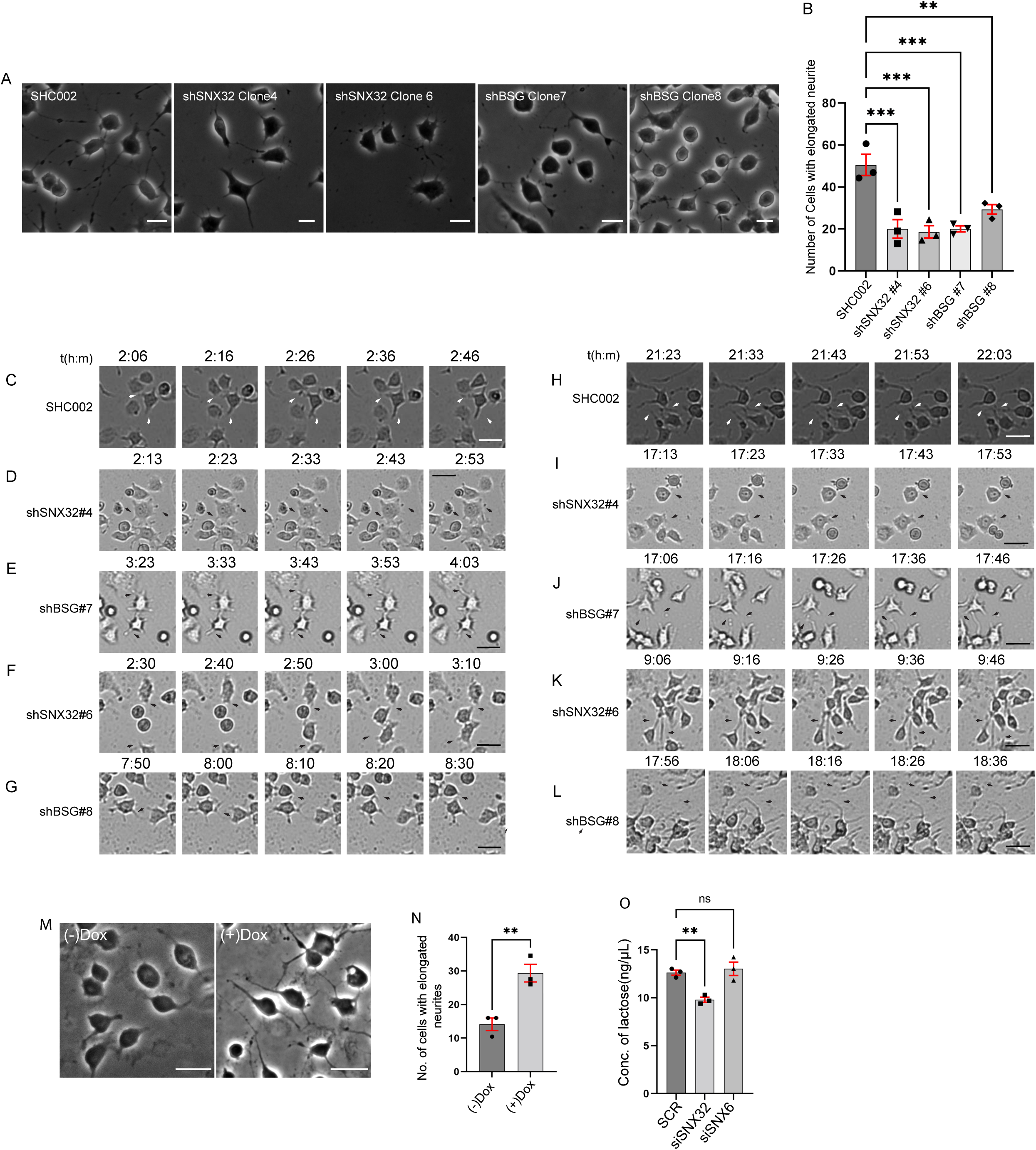
SNX32 and BSG plays a crucial role in neurite network formation: A) Phase contrast images of Neuro2a cells, which were transfected with Scrambled (SHC002)/ SNX32(shSNX32#4, shSNX32#6)/ BSG (shBSG#7, shBSG#8) shRNA clones followed by neurite induction as described in the materials and methods section, fixed and imaged using Zeiss Axio vert. A1 microscope, Scale Bar 50µm. B) Quantification of number of cells with elongated neurites, data represent mean ±SEM (N=3, Q≥100cen SaHU LQGHSHQGHQW H[SHULPHQWV, values are means ± SEM), P value 0.0003 (** P < 0.01, *** P < 0.001), Ordinary one-way ANOVA Dunnett’s multiple comparisons test. C-G) Snap Shots of Neuro2a cells transfected with Scrambled (SHC002)/ SNX32(shSNX32#4, shSNX32#6)/ BSG (shBSG#7, shBSG# 8) shRNA clones followed by neurite induction with RA for neurite induction showing neurite sprouting, Scale bar 50 µm (black arrows pointing to sprouting). H-L) Snap Shots of Neuro2a cells transfected with Scrambled (SHC002)/ SNX32(shSNX32#4, shSNX32#6)/ BSG (shBSG#7, shBSG# 8) shRNA clones followed by neurite induction with RA for neurite induction showing neurite extension, black arrows pointing network formation/retraction of neurites, Scale bar 50 µm. M) Phase contrast image of Neuro2a cells stably expressing pLVX SNX32 transfected with shSNX32#4 followed by neurite induction with or without doxycyclin as described in the materials and methods section, fixed and imaged using Zeiss Axio vert. A1 microscope, Scale Bar 50µm. N) Quantification of number of cells with elongated neurites, data represent mean ±SEM (N=3, n≥100cells per independent experiments), P Value 0.0091(** P < 0.01), Unpaired t test. O) Quantification of concentration of Lactate in the culture supernatant of U87MG cells transfected with SCR/SNX32/SNX6 SMARTpool siRNA, N=3, values are means ± SEM, P value 0.0049(** P < 0.01, ns-nonsignificant), Ordinary on e-way ANOVA Dunnett’s multiple comparisons test.

To validate our observations and further ensure the gene-specific effect, we performed a rescue experiment utilizing shRNA resistant SNX32 construct (pLVX shSNX32#4r). Neuro-2a cells stably expressing doxycycline-inducible pLVX shSNX32#4r was transfected with shSNX32#4 (SNX32KD) to acutely deplete endogenous SNX32. The SNX32KD cells were differentiated to form neurites by replacing the standard growth media (MEM with 10%FBS) with the differentiation media (MEM with 1%FBS containing 10 µmol/L Retinoic acid). The cells were allowed to grow in differentiation media with or without doxycycline for the next 48hrs. Further, the cells were fixed, and phase contrast images were acquired by AxioVert.A1 microscope. Neurite length was measured using ImageJ software and only neurites measuring more than the diameter of the cell body in length were taken for final analysis. It was observed that compared to no doxycycline-supplemented cells the doxycycline-supplemented cells showed an increased number of cells with elongated neurites (Fig. 7 M, N). To better understand and negate artefacts due to cell fixation, the procedure of neurite induction was replicated, and images were captured in real-time for a maximum of 48hrs (Supp. video 7). The DIC time-lapse live-cell video imaging showed that in doxycycline supplemented cells, long neurites became prevalent, but not in cells without doxycycline (Supp. video 7).

### SNX32 DOWNREGULATION DISRUPTS THE MCT - MEDIATED LACTATE SHUTTLING

Neuro-Glial coordination is necessary for the establishment of neuronal networks. The constant shuttling of metabolic fuels such as ketone bodies, pyruvate and lactate^45^ plays a major role in contributing to this coordination. The monocarboxylic acid transports (MCTs) play a critical role in Neuro-Glial coordination^46^ by facilitating the transport of monocarboxylates ^47^. Lactate is the most common substrate for MCT transport in the brain, where it is transported from its site of synthesis to the site of consumption between the glia and neurons^48^. Accordingly, the reduced lactate shuttle between Neuro-Glial cells interferes with the neurite outgrowth^49^. Therefore, we sought to quantify the extracellular concentration of lactate in cultured U87MG cells under SNX32 depleted condition.

We performed lactate quantification assay under SNX32KD or BSGKD conditions. The knockdown efficiency obtained by employing siRNA or shRNA in U87MG cells was measured at the mRNA level using qRT-PCR (Fig. S6 C-E). The culture supernatant of SNX32KD/BSG KD cells were collected and the amount of lactate was quantified following the manufacturer protocol. As reported earlier, BSG depletion led to reduced lactate concentration in the culture supernatant. We observed a similar reduction of lactate concentration in SNX32 downregulated condition (Fig. 7 O, Fig. S6 F). In contrast, the SNX6KD, the paralogue of SNX32, did not show any significant effect (Fig. 7 O). Our results suggest that in addition to its functional link with BSG, SNX32 is important for maintaining the activity of MCTs.

### SNX32 IS ESSENTIAL FOR THE SURFACE TRAFFICKING OF BSG

It has been reported that BSG acts as a cochaperone for MCTs and facilitates their surface localization^50^. Accordingly, BSG knockdown causes the accumulation of monocarboxylate transporters in the endo-lysosomal compartment, leading to reduced lactate concentration in the culture supernatant^51^. Therefore, we asked whether SNX32 regulates MCT activity via its role in BSG trafficking. When expressed from a transiently transfected inducible vector in U87MG cells, GFP-SNX32 showed considerable colocalization with BSG (Fig. S7 A). As previously reported, a significant population of BSG colocalized with ARF6 at the cell surface^52^ (Fig. 8 A, B). Similarly, in Neuro2a cells stably expressing cMyc-tagged BSG, there was considerable colocalization with GFP-SNX32 on intracellular punctae (Fig. 8 C), which was further confirmed by live-cell video microscopy (Supp. video 8). Next, we sought to assess if the surface population of BSG is altered in SNX32 downregulated conditions using TIRF microscopy. We utilized a Neuro2a cell line stably expressing an inducible vector pLVX TRE3G containing pHluorin^53^ tagged BSG. The SNX32KD/SNX6KD Neuro2a cells were supplemented with doxycycline to induce the expression of pHluorin-BSG, the cells were imaged in TIRF-M for a duration of 1min continuous timelapse without interval. The pHluorin-BSG punctate was identified in motion-tracking software and the number of vesicles was calculated per cells. The normalized number of vesicles was obtained by averaging the number of vesicles over total number of frames per cell. We observed that compared to scrambled (SCR) siRNA (Fig. 8 D), the ON-TARGETplus SMARTpool mediated downregulation of SNX32 (SNX32KD) in Neuro2a cells substantially reduced the surface population of pHluorin-BSG (Fig. 8 E, Supp. video 9) Moreover, we validated our observation using shRNA-mediated SNX32 downregulation (Fig. S7 B-E, Supp. video 10). Interestingly SNX6KD did not show any effect (Fig. 8 F, G), consistent with the absence of interaction between SNX6 and BSG. Furthermore, immunoblot analysis showed that in SNX32 silenced conditions, the protein levels of BSG were subtly reduced compared to Scrambled (SCR/SHC002) or SNX6KD conditions (Fig. S7 F-G). SNX32, therefore, regulates the cell surface trafficking of BSG, consistent with a working hypothesis that the role of SNX32 in neurite development is in part attributed to its ability to regulate the trafficking of BSG and, thereby, monocarboxylate transporters.

**Figure 8.**
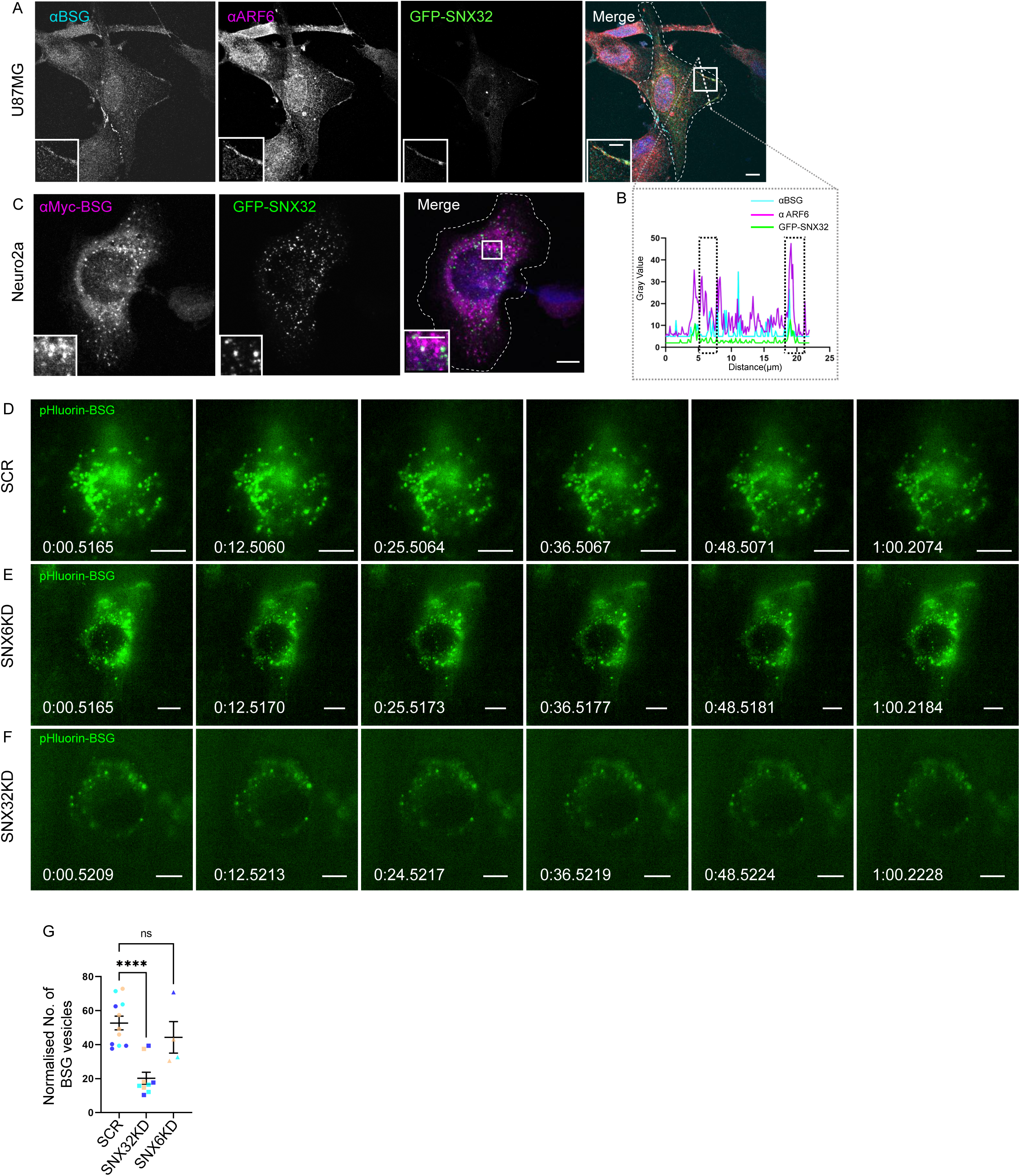
SNX32 but not SNX6 plays a significant role in surface localization of BSG: A) U87MG cells showing the colocalization of GFP-SNX32 with endogenous ARF6 and BSG on membrane, Scale 10µm, inset 5µm (magnified regions are shown as insets). B) Representative line intensity plot of U87MG cell showing intensity overlap of GFP-SNX32, ARF6 and BSG.C) Neuro2a cells showing colocalization of GFP SNX 32 with cMyc-BSG on vesicles, Scale 10µm, inset 5µm (magnified regions are shown as insets). D-F) Snap shots from live TIRF microscopic imaging of Neuro2a cells stably expressing pHluorin BSG transfected with SCR, SNX32 or SNX6 siRNA SMARTpool followed by doxycycline treatment for pHluorin BSG induction, Scale 10µm. G) Quantification of surface population of normalised of BSG vesicles, (N=3, n≥6cells per independent experiments), P value 0.0003(**** P < 0.0001, ns-nonsignificant), Ordinary one-way ANOVA Dunnett’s multiple comparisons test.

**Figure 9:**
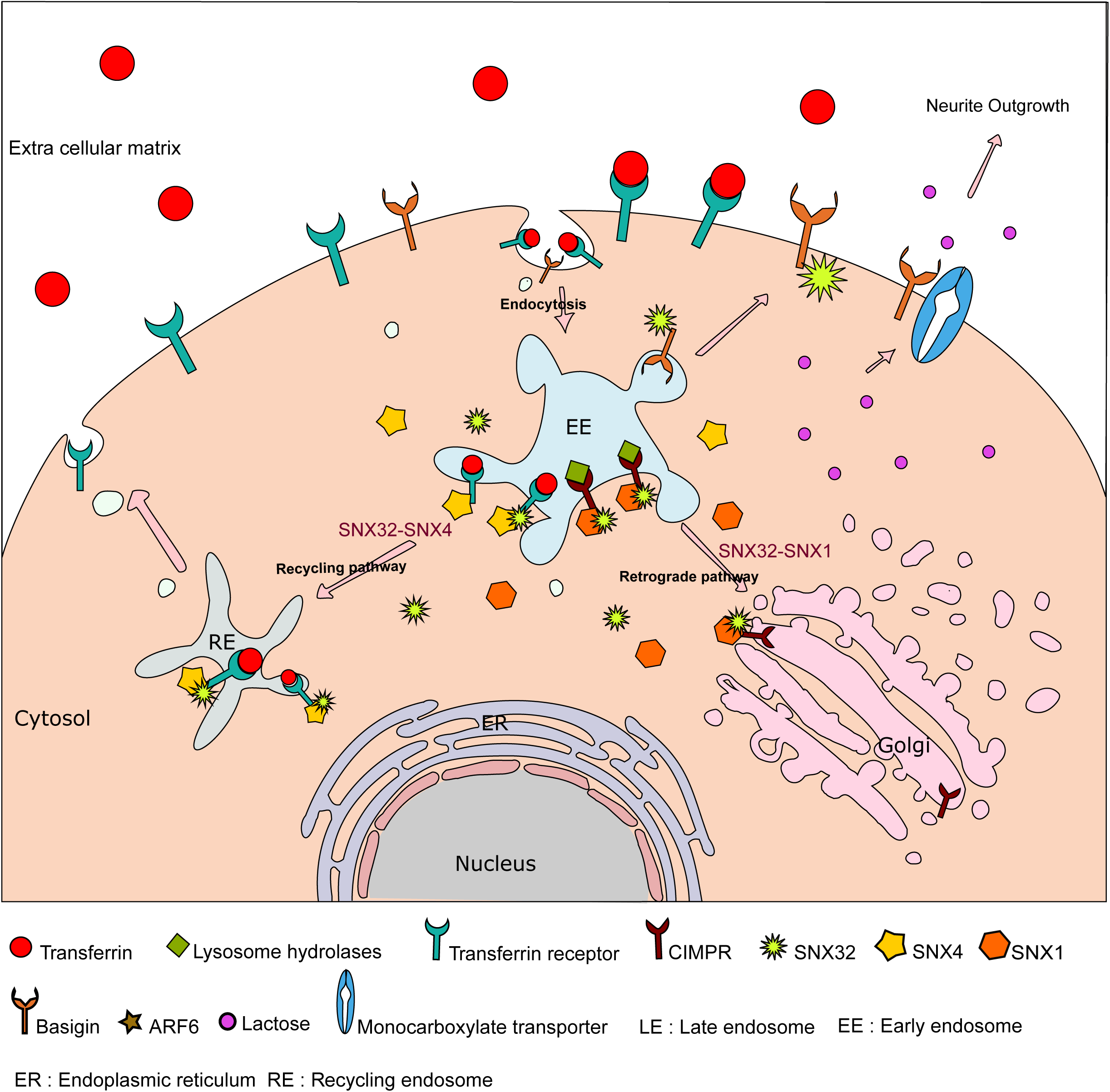
Graphical abstract showing SNX32’s multifaceted role in trafficking and Neuro-Glial coordination: A) SNX32 through its BAR domain associate with SNX4, co-traffic TfR from early endosome to recycling endosomes. Similarly, SNX32 interacts with SNX1, co-traffic CIMPR from early endosome to TGN. Further, the proposed role of SNX32 contributing to the endosome to surface trafficking of BSG. BSG being the co-chaperone of monocarboxylate transporter (MCT)^20, 26, 54^ helps in the surface localisation of MCTs. In conclusion, the derailing of SNX32 mediated BSG trafficking reduce the surface population of BSG as well as MCTs, disrupting the Neuro-Glial coordination and manifests as neurite differentiation defect *in cellulo.* The model depicts the diverse cargo trafficking route in which SNX32 functions, and a glimpse of complexity introduced by means of its ability to participate into distinct protein complexes.

## DISCUSSION

### Localization of SNX32 is assisted by protein-protein interactions

The cargo sorting at endosomal compartments is a multi-stage process comprising membrane recruitment of the cargo identification complex, recognition of cargoes, membrane tubulation, vesicle fission, and cargo dislodging at the target membrane. The evolutionarily conserved SNX-BAR family of proteins are implicated in cargo identification, sorting, and membrane tubule biogenesis^54, 55^. The co-dependence of SNXs owing to the ability of SNX-BARs to undergo homo/hetero-dimerization is a critical feature during various stages of cargo sorting ^10, 25, 32, 56, 57^. In HeLa cells, our results indicate that the BAR domain-mediated interaction with SNX1/SNX4 might contribute to the recruitment of SNX32 to EEA1, harbouring early endosomes (Fig.2a, Fig.S2a), also supported by the SNX32ΔN’s localization at early endosome and its sensitivity to PI3K inhibitor-wortmannin. Though earlier reports have shown that SNX32, similar to its closest homologs SNX5 and SNX6, was unable to induce *in vitro* membrane tubulation^2^, our results indicate that it may not be due to the SNX’s insufficiency in membrane association. Based on our results, we hypothesize that SNX32 engages in a heteromeric interaction with a partner, efficient in inducing membrane tubules such as SNX4, while it itself contributes to cargo recognition through its PX domain (Fig.). Similar interdependence was also observed for SNX6 and SNX27, where their hetero-dimerization with SNX1/2 or SNX1, respectively, was shown to be important for membrane recruitment ^57, 58^. Further, it is noted in heterodimeric associations of SNX1/SNX5 and SNX1/SNX6, while SNX5, SNX6 lead cargo recognition^1, 10, 11, 25^, SNX1, SNX2 are known to contribute to membrane remodelling establishing the mutual cooperativity of the individual SNX proteins during cargo sorting.

The hydrophobic interactions are important for the stabilizations of BAR domain dimerization^2^. In addition, a few polar interactions, including hydrogen bonds and salt bridges, also aids in defining the specificity in complex formations^59^. A similar trend is also observed for SNX32-SNX1 and SNX32-SNX4; the interface is largely dominated by hydrophobic interactions along with multiple polar interactions (Fig. 2 c,f). In the current study, we were also able to identify A226, Q259, E256, R366 of SNX32, and Y258, S448 of SNX4, which were present in the SNX4-SNX32 interface, and mutating these residues, which contribute to the stabilization of the heterodimeric interface between SNX4 and SNX32 could indeed abrogate the interactions. It will be interesting to see the physiological implications of these mutants. Additionally, we also observed that SNX32 interacts with the other SNX-BAR family member, SNX8 (Fig. 1a). Keeping in an account SNX8’s role in early endosome to TGN transport ^60^ it will be interesting to explore whether/how this two SNXs pair-up to facilitate cargo sorting.

### The lipid binding and cargo recognition ability of the PX domain accounts for its role in cargo sorting

Though the contribution of heteromeric interactions in membrane localization of SNX32 is in place, the role of PIns affinity of the PX domain cannot be disregarded. The purified PX domain of SNX32 showed preferential binding to PI3P in our lipid overlay assay (Fig1.j), which along with the results from our cellular localization study on SNX32-PX in the presence or absence of Wortmannin (Fig.S1m), indicated that SNX32’s PX domain might also contribute to its localization on early endosomes. Likewise, its affinity towards PI4P, as evident from (Fig1.j) along with its localization in PAO-treated cells (Fig. 1l), corroborates well with SNX32’s association with Golgi/recycling compartments. Given the presence of PI4P in the microdomains of the cell surface, the observed localization of SNX32 in HeLa/Neuro/Glia (Fig. 3i-j, Fig.1d) and the functioning of the SNX32 in these cell lines are not surprising. The lipid affinity of the purified PX domain of SNX32(Fig1.j) indicates that the differential phospholipid affinity of the complex might be playing a role in leading the cargo through endosomal maturation processes. In this context, the role of motor proteins in driving the cargo containing vesicles from the source compartment to the destination is of particular interest. For instance, in the case of SNX1/5 and SNX1/6 complex, the interaction of SNX5/SNX6 with p150glued, an activator of motor protein Dynein, is necessary for the tubular sorting carrier formation^57, 61^. Further, the affinity difference between the interaction with the motor and the PIns in the destination membrane commission the cargo dislodging, as delineated in the case of SNX1/SNX6 complex^25, 57^. Since SNX32 belongs to the SNX5/SNX6 cluster, the possibility of SNX32 carrying out the cargo sorting in a similar course is highly probable (Fig. 4a). In addition, the investigation of CIMPR/TfR trafficking was focused on understanding the contribution of SNX32 in retrograde/recycling trafficking route.

In our study, we have observed that SNX32 is able to interact with multiple cargoes as well as with multiple SNX-BAR family members. We were able to identify the F131 of SNX32 as a crucial amino acid in mediating the interaction with TfR, CIMPR and BSG. Though the mechanistic insight into the sorting events is still enigmatic, it is likely the drill would be similar to that observed for β2-adrenergic receptor or Wntless^62^ where the sorting is ultimately limited/decided by the relative concentration of the cargoes in membrane sub domains^1^.

Briefly, in the current study, we showed that SNX32, an SNX-BAR protein shown to be insufficient in inducing membrane remodelling^2^, through its BAR domain interacts with SNX1 or SNX4, which show an intrinsic membrane remodelling characteristic^30^. Once recruited to the endosomal membrane, it’s the PX domain of SNX32 that interacts with the cargo, possibly assisting in the assemblage leading to cargo-enriched subdomains formations within the sorting endosome. Based on the earlier report ^25^ it could be hypothesized that, once the vesicles are dispatched to the destination, it’s the affinity of the SNX32 PX domain with PI4P that make for cargo dislodgement at the destination. It will therefore be interesting to delineate the mechanism underlying the selective nature of cargo recognition by distinct heteromeric complexes of SNX proteins.

### SNX32-mediated sorting critically contributes to neuronal differentiation

The role of SNX in brain functionality and neuronal disorder is an emerging area of research^7, 63^. The contribution of multiple sorting nexins in amyloid precursor protein (APP) trafficking and metabolism have been actively investigated in recent years^7, 64–66^. In this context, it is interesting to note that the interactome of SNX32 encompasses proteins such as PGRMC1^38^, ROBO1^39, 67–69^, SV2a^40^ and BSG, which are known to be vital in neuronal development^10^. The current study begins to delineate a role for SNX32 in regulating neuronal differentiation through its cargo protein, BSG. BSG contributes to a plethora of physiological functions, including sensory and nervous system functioning^41, 70, 71^. BSG’s interaction with integrin is necessary for cytoskeletal rearrangements^72^, which is crucial during neurite outgrowth. Our data suggested that SNX32 regulates the surface localization of BSG(Fig) via its role in plasma membrane recycling of the later from endosomes, explaining the observed phenocopying of SNX32 and BSG. Further, it has been shown that BSG acts as a chaperone for the monocarboxylate transports (MCTs), which plays a crucial role in lactate shuttling and thereby contributes to energy metabolism in neurons. Since lactate accounts for a significant share of energy sources in neurons, the reduced lactate shuttle consequently contributes to defects in neuronal development^46^. Thus, SNX32, via its role in cell surface transport of BSG, may also regulate MCTs and consequently contribute to energy metabolism in developing neurons (Fig. 6). Moreover, the enrichment of kinesin-1 heavy chain (KiF5b) in the differential proteomics data of SNX32 suggests that this motor protein may be a potential interactor of SNX32 in neuronal cells. Being a plus-end directed motor, KiF5 plays an essential role in cargo transport during neurite extension, and accordingly, inhibition of KiF5 abrogates axon specification^35^. Taken together, it is tempting to speculate that the association with KiF5, may enable SNX32 to contribute to the long-range trafficking of proteins like BSG to the surface.

## Materials and Methods

### Plasmids and antibodies

pEGFP C1 SNX32 FL, pEGFP C1 SNX1, pEGFP C1 SNX4, mCherry C1 SNX1 where a kind Gift from Prof Peter J Cullen. pGEX6P1 GFP Nanobody (#61838, Addgene) was purchased from Addgene and subcloned to pGEX6P1 for improved induction, pGEX6P1mCherry nanobody (#70696, Addgene) was purchased from Addgene. pLVX TRE 3G and pLVX EF1Į Tet3G were part of Tet-On® 3G Inducible Expression Systems (631363, Takara Bio Inc.). pCMV6-Entry Myc DDK CD147 (BSG) (#: RC203894 Origene). pLVX TRE3G pHluorin BSG was subcloned from #RC203894, pLVX-TRE3G GFP-SNX32#4r was synthesised by commercial cloning service provider GenScript USA Inc. pmCherry-Rab 11 was a kind Gift from Prof. Marino Zerial. pIRES neo2 CD8 CIMPR was a kind Gift from Prof. Matthew N J Seaman. pmCherry C2-PH ^PLCδ^ was a kind Gift from Prof. Pietro De Camilli. pEGFP C1 PH ^OSBP^ was a kind Gift from Prof. Tamas Balla. pLVX TRE3G GFP-SNX32FL, pcDNA3 HA N(I) SNX32 FL, pmCherry-C2 SNX32 FL, pcDNA3 HA N(I) SNX 32 ΔN, pcDNA3 HA N(I)SNX32 ΔC, His-SNX32FL, His-SNX32ΔN, His-SNX32ΔC, GST-SNX32FL, GST SNX32ΔC where subcloned from pEGFP C1 SNX32FL, RFP C1 SNX4 was subcloned from pEGFP C1 SNX1. pcDNA3 HA N(I) SNX32#4r, pcDNA3 HA N(I) SNX32#4rΔN, pEGFP SNX4Y258E, pEGFP SNX4S448R, pcDNA3 HA N(I) SNX32 R220E, pcDNA3 HA N(I) SNX32A226E, pcDNA3 HA N(I) SNX32E256R, pcDNA3 HA N(I) SNX32 Q259R, pcDNA3 HA N(I) SNX32 R366E were constructed using site directed mutagenesis.

Antibodies used in this study are anti-GFP (11814460001, Roche; WB,1:3000), anti-HA(C29F4, Cell Signalling Technology,WB,1:1000, IF,1:500),anti-His (MA1-21315, Invitrogen; WB,1:10,000), anti-EEA1 antibody was a kind gift from Prof. Marino Zerial, anti-HA(sc7392, Santa Cruz, WB/IF, 1:500),anti-TGN46(AHP1586,AbD Serotec, IF,1:200), anti-transferrin receptor (13-6800, Invitrogen; WB,1:1000, IF,1:200), Transferrin 488(T13342, Invitrogen, IF,5µg/ml), Transferrin 568(T23365,Invitrogen,IF,5µg/ml), Transferrin 647(T23366 Invitrogen, live imaging, 10µg/ml),anti-SNX1(611482,BD Biosciences,IF,1:500) anti-Vinculin (V9131, Sigma Aldrich, WB,1:1000), anti-CIMPR (ab2733, Abcam,IF,1:500), anti-CD8(153-020,Ancell,IF, 1:500, Live imaging, 1:300), anti-CIMPR (ab124767, Abcam,WB,1:50,000), anti-mCherry (M11217, Invitrogen; WB, 1:1000,IF, 1:500), anti-CD147(345600, Invitrogen; WB/IF, 2µg/ml),c-Myc (9E10-sc40, Santa Cruz,IF,1:500),anti-ARF6 antibody was a kind gift from Dr.Vimlesh Kumar.

HeLa cells were a kind gift from Prof. Marino Zerial, Max Planck Institute of Molecular Cell Biology and Genetics, Dresden. U87MG and Neuro2a cells were acquired from Cell Repository, National Centre for Cell Science Pune, India.

### Uptake media composition

**For HeLa**–DMEM containing 10% heat-inactivated South American FBS, 1x Penicillin/Streptomycin, 20mM HEPES (pH 7.5)

**For Neuro2a**-MEM containing 10% heat-inactivated South American FBS, 1x Pencillin/Streptomycin,2mM Sodium Pyruvate, 20mM HEPES (pH 7.5)

### Cleared cell lysate preparation

Cells were lysed with Lysis buffer (50mM Tris (pH 8),150mM NaCl,0.5% NP40) for 5min on rotamer. Later, the lysates were cleared by centrifugation at 19000g for 20min at 4°C. The supernatant fraction was collected without disturbing the pellet and used for experiments.

### siRNA transfection

All ON-TARGET plus siRNA SMARTpools were purchased from Dharmacon. HeLa cells were transfected with SMARTpool siRNAs against negative control siRNA-Scrambled(D-001810-10), SNX32 (L-017082-01), SNX6 (L-017557-00). Cells were transfected with 15–20 nm siRNA using DharmaFECT 1 (Dharmacon) following manufacturer protocol and incubated for a maximum of 72 hrs before any further analysis.

### shRNA Transfection

All shRNA clones were part of the MISSION shRNA product line from Sigma Aldrich. The TRC1.5 pLKO.1-puro non-Mammalian shRNA Control Plasmid DNA (SHC002) is a negative control containing a sequence that should not target any known mammalian genes but engage with RISC. The SNX32 Clones – shSNX32#4 (TRCN0000181072), shSNX32#6 (TRCN0000180862), and the BSG Clones-shBSG#7 (TRCN0000006733), shBSG#8 (TRCN0000006734) were screened for maximum knockdown efficiency compared to other available clones targeting the same protein. The cells were transfected with shRNA clones (as mentioned in the figures and legends) using Lipofectamine LTX with Plus Reagent (Invitrogen) for a maximum of 48hrs before any further analysis.

### shRNA resistant SNX32

pLVX TRE 3G SNX32 resistant (shSNX32#4r) to shRNA clone – shSNX32#4 (TRCN0000181072) was synthesised by commercial cloning service provider GenScript USA Inc., which was further subcloned into pcDNA3 HA vector backbone. pcDNA3 SNX32ΔN resistant (shSNX32ΔN#4r) was constructed following site directed mutagenesis using the primer 5’ TTCGAACACGAACGGACAT 3’.

### GFP/mCherry nanobody mediated immunoprecipitation

20 µg of GST-GFP/mCherry nanobody was allowed to bind to glutathione Sepharose beads in 1X PBS for 1 h at 4°C. Followed by removal of unbound proteins, the beads were incubated with cell lysate containing overexpressed GFP /mCherry empty vector, or GFP/mCherry tagged target protein for 2hrs at 4°C. After incubation, unbound protein residues were removed by washing the beads-nanobody-Target protein complex thrice with 400µl 1xPBS. Further, samples were prepared, resolved on SDS-glycine gel, and analysed after immunoblotting. 5% of the cleared cell lysate was used as input.

### Site directed mutagenesis

Mutagenesis PCR was used to introduce mutations into SNX4 and SNX32. A mutant construct was generated by PCR, using GFP-tagged SNX4 or HA-tagged SNX32 as template DNA, with an appropriate forward primer and a reverse primer that introduced the mutation. The PCR product was treated with Dpn1 to digest the template and hybrid DNA, followed by transformation to DH5Įcells. The constructs confirmed by DNA sequencing.

### Indirect immunofluorescence

Cells were fixed using 4% Paraformaldehyde (PFA) or 100% methanol (MeOH, Sigma Aldrich) for 10mins (PFA) or 15mins (MeOH) at room temperature (PFA) or −20°C(MeOH). Following PFA fixation, the cells were permeabilized using 0.1% triton-x-100(Sigma Aldrich) in PBS. Cells fixed using MeOH were directly blocked using 5% Foetal Bovine Serum in PBS before incubating with primary and secondary Alexa labelled antibodies. The coverslips (1943-10012A, Bellco) were mounted using Mowiol on glass slides and imaged using Zeiss LSM 780 laser-scanning confocal microscope 63 × /1.4 NA oil immersion objective lens or Olympus FV3000 confocal laser-scanning microscope with a 60 × Plan Apo N objective (oil, 1.42 NA). Data from three independent experiments were subjected to analysis by the automated image analysis program, Motion Tracking (http://motiontracking.mpi-cbg.de). The imaging frames were randomly selected, and a minimum of 15 images were acquired for each experimental condition in a given setup. Further, all the images were pooled and processed together. The objects were identified based on their size, fluorescence intensity, and other parameters by Motion Tracking software. Objects detected in two different channels were considered colocalized if the relative overlap of respective areas was >35%. The apparent colocalization value was calculated and corrected for random colocalization.

### Phenyl Arsenide Oxide/Wortmannin treatment

Phenyl arsenide oxide /Wortmannin treatment was followed as previously reported^57^. Briefly, HeLa cells were transiently transfected with individual target protein plasmids and incubated for 12-14hrs at 37°C, 5% CO_2_. Further, the cells were treated with PAO (15µM)/ Wortmannin (200nM) in uptake medium for 15mins at 37°C, proceeded for Immunofluorescence.

For the *in vitro* membrane relocalization assay, following the PAO treatment, cells were used for membrane fractionation assay and equal amount of protein was resolved using SDS-PAGE and analysed after immunoblotting.

### Generation of stable cell lines

HeLa cells stably expressing GFP-SNX32 from an inducible promoter were established following the distributor protocol (Takara Bio Inc.). GFP-SNX32 full-length gene was cloned onto pLVX-TRE3G Vector. For viral titer preparation, 7µg of pLVX-TRE3G GFP-SNX32 or pLVX-pEF1a-Tet3G Vector plasmid DNA was diluted in sterile water to a final volume of 600 µl. Further, the diluted DNA was added to Lenti-X Packaging Single Shots for 10 min at room temperature to allow nanoparticle complexes to form. Followed by incubation, the entire 600 µl of nanoparticle complex solution was added to 80% confluent HEK-293T cells. The cells were incubated at 37°C, 5% CO_2_ for 24–48 hrs. Viral titers were harvested by centrifuging briefly (500g for 10 min) followed by filtering through a 0.45-µm filter to remove cellular debris. Later, HeLa cells were transduced with the viral titers of pLVX-TRE3G GFP-SNX32 and pLVX-pEF1a-Tet3G mixed in 1:1 ratio and topped up with DMEM complete media containing 4 µg/ml concentration of Polybrene. 24 hrs past transduction, the culture media was removed and replaced with complete DMEM containing G418(600µg/ml) and Puromycin (10 µg/ml) selection media. Allowed the cells to divide and form colonies for 14days before passaging. Cells were maintained in complete DMEM containing G418 (400µg/ml) and Puromycin (0.25 µg/ml).

Neuro2a cells stably expressing pHluorin BSG from an inducible promoter were established by co-transfecting pLVX-TRE3G pHluorin BSG, pLVX-pEF1a-Tet3G Vector plasmid DNA in 1:1 ratio using Lipofectamine LTX with Plus Reagent (Invitrogen). 24hrs post-transfection, the culture media was replaced with complete MEM containing G418(600µg/ml) and Puromycin (10 µg/ml) selection media. Allowed the cells to divide and form colonies for 18days before passaging. Cells were maintained in complete MEM containing G418 (400µg/ml) and Puromycin (0.25 µg/ml).

### Lipid overlay assay

Lipid overlay assay was performed using PIP strips (nitrocellulose membrane spotted with 8 phosphoinositides and 7 other biologically relevant lipids) according to the manufacturer’s instructions (Thermo Scientific). Briefly, the membrane was incubated with 0.7 mg/ml His SNX32ΔC diluted in blocking buffer (TBS-T + 3% BSA) for 13hrs at 4°C with gentle mixing. The His SNX32ΔC protein bound to the lipids was detected by anti-His antibody [1:10,000 (Thermo Scientific)]

### SIM sample preparation

As mentioned earlier, HeLa cells expressing the protein of interest were fixed in 4% Paraformaldehyde (PFA) for 10min at room temperature and proceeded for indirect immunofluorescence. Briefly, the cells were permeabilized with 0.1% Triton X 100 for 12 minutes at room temperature and immunostained using GFP, mCherry primary antibodies, followed by Alexa labeled secondary antibodies. The coverslips were mounted on glass slides using mounting media without DAPI. Image captured using Nikon N-SIM S system in 3D-SIM mode (sequentially) with laser wavelengths 488nm and 561nm. For each Z plane and for each wavelength 15 images were captured (3 different angle and 5 different phases). Images were captured in Nikon N-SIM S demo system and reconstructed using Nikon software NIS Elements version 5.30.

### Membrane fractionation and enrichment assay

The assay was performed as previously reported^29^.Briefly, Hela cells grown in T25 tissue culture flasks were washed twice with PBS and drained completely. Later, the cells were snap frozen using liquid nitrogen and then quickly thawed at room temperature. Cells were scraped off in 0.5 ml of lysis buffer (0.1 M Mes-NaOH pH 6.5, 1 mM magnesium acetate, 0.5 mM EGTA, 0.2 M sucrose and 1xPIC). The pellet (which contained the membrane proteins) was separated from the supernatant (which contained the cytosolic proteins) by centrifuging at 10000xg for 10min. The pellet was then solubilised in 0.3 ml of lysis buffer (50 mM Tris-HCl, pH 7.4, 150 mM NaCl, 1 mM EDTA, 1% Triton X-100, 0.1% SDS) before next round of centrifugation at 10,000 × *g* for 10 minutes. The supernatant from the second spin now contained membrane and membrane-associated proteins, which was used as membrane-enriched fraction for *in vitro* pulldown assay. Equal amount of pellet and supernatant was loaded for analysing membrane re-localization assay in the presence of PAO.

### Transferrin pulse-chase assay

This was followed as previously reported ^73^. Briefly, HeLa cells were transfected with scrambled/ individual siRNA SMART pool or shc002/shSNX32 clones for 70hrs(siRNA) or 46 hrs (shRNA) followed by serum starvation for 2 hours to deplete the endogenous transferrin population. Cells were then washed with uptake media and incubated with 5µg/ml transferrin (Alexa Fluor 488/568 conjugated) at 37°C for 30 minutes. After completing the pulse period, the cells were washed and proceeded for chase using unlabelled Holotransferrin 100µg/ml. Cells were fixed at specified periods and proceeded for immunofluorescence.

### CD8uptake assay

HeLa cells stably expressing GFP-Golph3 and CD8-CIMPR were transfected with scrambled/ individual siRNA SMART pool or shc002/shSNX32 clones for 72hrs(siRNA) or 48 hrs (shRNA). Further, cells were incubated with anti-CD8 monoclonal antibody for 60 minutes on ice, followed by two quick washes with uptake media to remove unbound antibodies. Chase was done by incubating the cells in pre-warmed uptake media at 37°C for 30mins followed by fixation and proceeded for immunofluorescence.

### Protein purification

#### Purification of His SNX32ΔC

E. coli BL21 (DE3) cells were transformed with plasmids encoding His SNX32ΔC. Colonies were screened, and culture induction conditions were standardized. The culture was grown at 37 °C until OD600 reached ~0.6–0.8. Temperature was then lowered to 16 °C, and protein expression was induced by adding 0.1 mM isopropyl β-d-1-thiogalactopyranoside (IPTG). After ~15 h, cells were harvested and homogenized in lysis buffer (20 mM Tris, pH 8.0, 400 mM NaCl, 2 mM 2-Mercaptoethanol (βME), 10 mM Imidazole and 1 mM PMSF). Cells were lysed by sonication for 2min and subjected to high-speed centrifugation to remove insoluble debris. The supernatant was then mixed with Ni-NTA beads for 20 min. Unbound proteins were washed off with wash buffer (20 mM Tris (pH 8.0), 400 mM NaCl, 2 mM βME, 10 mM Imidazole). Protein was eluted in buffer containing 250mM Imidazole. The eluted protein was subjected to buffer exchange (20 mM Tris (pH 8.0), 400 mM NaCl, 2 mM 2-Mercaptoethanol (βME),10%glycerol) in Jumbosep™ Centrifugal Devices and flash frozen to store at −80°C.

#### Co purification of GST SNX1/ His SNX32ΔN

The procedure was followed as reported by Yong et al.,^74^. Briefly, plasmids encoding GSTSNX1 and His SNX32ΔN were co-transformed into E. coli BL21 (DE3) strain, grown in Luria-Bertani (LB) agar plates supplied with 50 μg/ml ampicillin and 30 μg/ml kanamycin. An overnight liquid culture of 10ml was used to initiate a 1L expression of SNX1/SNX32ΔN complex. The culture was grown at 37 °C until OD600 reached ~0.6–0.8, later the temperature was lowered to 22 °C, and protein expression was then induced by adding 0.5 mM isopropyl β-d-1-thiogalactopyranoside (IPTG). After 14-15 h, cells were harvested and homogenized in lysis buffer (20 mM Tris, pH 8.0, 400 mM NaCl, 2 mM 2-Mercaptoethanol (βME), 10 mM Imidazole, 10% (v/v) glycerol, and 1 mM PMSF). Cells were lysed by sonication for 2min (10sec ON-10sec OFF Pulses) and subjected to high-speed centrifugation to remove insoluble debris. The supernatant was then mixed with Ni-NTA beads for 20 min. Unbound proteins were washed off with wash buffer (20 mM Tris, pH 8.0, 400 mM NaCl, 2 mM βME, 10 mM Imidazole). Fractions were concentrated and flash frozen before storing at −80 °C.

#### SILAC

For SILAC, SH-SY-5Y cells stably expressing GFP or a GFP-tagged construct of the protein of interest were cultured for at least six doublings in SILAC DMEM (89985; Thermo Fisher Scientific) supplemented with 10% dialyzed FBS (F0392; Sigma-Aldrich). Cells expressing GFP were grown in media containing light amino acids (R0K0), whereas cells expressing the GFP-tagged protein of interest were grown in medium (R10K8 or R6K4). Amino acids R10,R6, R0, and K0 were obtained from Sigma-Aldrich, whereas K4 was from Thermo Fisher Scientific. Cells where lysed in immunoprecipitation buffer (50 mM Tris-HCl, 0.5% NP-40, and Roche protease inhibitor cocktail) and subjected to GFP trap (ChromoTek). Precipitates were pooled and separated on NuPAGE 4–12% precast gels (Invitrogen) before liquid chromatography–tandem mass spectrometry analysis on an Orbitrap Velos mass spectrometer (Thermo Fisher Scientific).

#### *In vitro* pulldown assay

20 μg of His SNX32ΔC was allowed to bind to Ni-NTA beads for 20min, 4°C with end-to-end mixing. The unbound protein was washed off with PBS, followed by incubation with membrane enriched HeLa cell lysate for 2hrs, 4°C with end-to-end mixing. The samples were resolved using SDS-PAGE and analysed after immunoblotting.

#### Neurite outgrowth assay

Neurite outgrowth assay was performed as previously reported^36^. Briefly, Neuro2a cells were transfected with Scrambled or individual siRNA SMART pools or shc002/shSNX32/shBSG clones. Following 24hrs(siRNA) or 6hrs(shRNA) of transfection, the medium was replaced with MEM containing 1% fetal bovine serum supplemented with 10 μmol/L retinoic acid (RA) for another 48hrs to induce neurite outgrowth. The formation of neurites was observed using Axio Vert.A1 Inverted Transmitted Light Microscope (Carl Zeiss Microscopy GmbH, Göttingen, Germany) after cell fixation or live images were captured using JuLI™Br inverted microscope (NanoEnTek). The JuLI™Br system is equipped with a station unit that runs inside a CO_2_-regulated incubator and a scope unit that runs outside the incubator. Phase-contrast images were captured every 1 min for 48hrs using a 4× objective and a CMOS camera with a pixel length of 0.586 µm. The cells with neurites were counted using Image J software.

### Rescue experiments

#### A) Transferrin Pulse-Chase

HeLa cells were transiently transfected with SNX32 shRNA clone shSNX32 #4 to deplete the expression of SNX32 for 34hrs. Further, the cells were transfected with shSNX32 #4 resistant HA tagged SNX32 for 12hrs; following that, the cells were serum starved for 2hrs. Transferrin Pulse-chase was carried out. Cells were fixed at specified periods and proceeded for immunofluorescence

#### B) Neurite outgrowth

Neuro2a cells stably expressing doxycycline-inducible pLVX SNX32#4 resistant construct was transiently transfected with shSNX32#4 to deplete the expression of SNX32 for 48hrs (until the completion of the experiment). Following 6hrs after transfection, the cells were induced with Doxycycline for expressing shSNX32 #4 resistant SNX32 and RA in reduced serum-containing media. The formation of neurites was observed using Axio Vert.A1 Inverted Transmitted Light Microscope (Carl Zeiss Microscopy GmbH, Göttingen, Germany) after cell fixation or live images were captured using JuLI™Br inverted microscope (NanoEnTek). The JuLI™Br system is equipped with a station unit that runs inside a CO_2_-regulated incubator and a scope unit that runs outside the incubator. Phase-contrast images were captured every 1 min for 48hrs using a 4× objective and a CMOS camera with a pixel length of 0.586 µm. The cells with neurites were counted using Image J software.

### Confocal live-cell microscopy

HeLa cells seeded on glass-bottom dishes were transfected with respective constructs using Lipofectamine LTX with Plus reagent (Invitrogen) and incubated for 12hrs, 37°C, 5%CO_2_. Images were acquired using Olympus FV3000 confocal laser-scanning microscope for supplementary videos 1 and 2.

To capture transferrin co-traffic, HeLa cells were transiently transfected with mCherry-SNX32 and GFP-SNX4. Later cells were serum-starved for 2hrs and incubated with Transferrin (Alexa Fluor 647 conjugated) in uptake media, for 2mins prior to imaging. Videos were captured for 5min, without intervals in Olympus FV3000 confocal laser-scanning microscope with a 60× Plan Apo N objective (oil, 1.42 NA) on an inverted stage. Images were acquired and processed using FV31S-SW software and ImageJ software, respectively.

To capture CD8-CIMPR co-traffic, HeLa cells stably expressing pLVX TRE3G GFP-SNX32 were transiently transfected with mCherry-SNX1 for 4hrs followed by doxycycline induction for 12hrs. Later, cells were treated with anti-CD8 antibody and incubated for 1hr in 4°C. Unbound antibody was washed using uptake media followed by incubation with Alexa 647 labelled secondary antibody. Cells were washed twice with uptake media prior to imaging. Videos were captured for 5min, without intervals in Olympus FV3000 confocal laser-scanning microscope with a 60× Plan Apo N objective (oil, 1.42 NA) on an inverted stage. Images were acquired and processed using FV31S-SW software and ImageJ software, respectively.

To capture GFP-SNX32 and Myc-BSG co-trafficking, Neuro2a cells were transiently transfected with pLVX TRE3g GFP-SNX32 and Myc-BSG for 4hrs followed by doxycycline induction for 12hrs. Later, cells were treated with anti-Myc antibody and incubated for 1hr in 4°C. Unbound antibody was washed using uptake media followed by incubation with Alexa 647 labelled secondary antibody. Cells were washed twice with uptake media prior to imaging. Videos were captured for 5min, without intervals in Olympus FV3000 confocal laser-scanning microscope with a 60× Plan Apo N objective (oil, 1.42 NA) on an inverted stage. Images were acquired and processed using FV31S-SW software and ImageJ software, respectively.

### Lactate quantification assay

Lactate quantification assay was performed following the manufacturer protocol (MAK017, Sigma Aldrich). Briefly, U87MG cells were plated on a 96-well plate (1000 cells per cell well) and cultured overnight in MEM containing 10%FBS, followed by treatment with scrambled/SNX6/ SNX32 siRNA SMART pools or SHC002/shSNX32/shBSG clones. After 72hrs(siRNA) or 48hrs(shRNA) of transfection, the supernatant medium was collected and used for Lactate quantification.

### TIRF microscopy

TIRF microscopy was performed using a Nikon Eclipse Ti2 microscope, equipped with an incubation chamber (37 °C,5% CO_2_), a ×60 TIRF objective (oil-immersion, Nikon), an sCMOS camera (Neo, Andor), a 100 W mercury lamp (C-LHG1 Mercury). Stable Neuro2a cells expressing pHluorin BSG were seeded on a glass-bottom dish (100350, SPL), followed by treatment with scrambled/SNX6/ SNX32 siRNA SMART pools or shc002/shSNX32 clones for a maximum of 72hrs(siRNA) or 48hrs(shRNA). 12hrs prior to imaging, cells were transfected with doxycycline for pHluorin BSG induction. Data from three independent experiments were subjected to analysis by the automated image analysis program, Motion Tracking (http://motiontracking.mpi-cbg.de). The imaging frames were randomly selected, and a minimum of 5videos of 1min duration, without interval, were acquired for each experimental condition in a given setup. The objects were identified based on their size, fluorescence intensity, and other parameters by Motion Tracking software. The number of objects detected was normalized with the size of the vesicles and averaged with the number of cells per frame were plotted and compared in GraphPad Prism 9.

## Acknowledgments

We extend our gratitude to Prof. Peter J Cullen and Dr. Boris Simonetti (University of Bristol, UK) for the fruitful discussions and the guidance they offered, and also providing the SILAC data. We also acknowledge Ms. Katy (Prof. Cullen’s group, University of Bristol) for resource sharing. We sincerely that Prof. Matthew N J Seaman (Cambridge Institute for Medical Research) for his critical comments on the study and also for resource sharing. We thank Prof. Marino Zerial (Max Planck Institute of Molecular Cell Biology and Genetics), Prof. Pietro De Camilli (Yale’s Boyer Center for Molecular Medicine), Prof. Tamas Balla (National Institutes of Health) for sharing plasmids for mammalian expression. We acknowledge the FIST facility at IISER Bhopal by DST for providing confocal microscopy facilities. We are grateful to Dr. Vimlesh Kumar (IISER Bhopal) for suggestions and for sharing the ARF6 antibody. We sincerely thank Dr. Raghuvir Singh Tomar, Dr. Vikas Jain, and Dr. Himanshu Kumar for providing access to various instruments. We thank Prabal Kumar Chakraborty and Suparno Gupta of Towa Optics (I) Pvt. Ltd. for assistance in image acquisition using the Nikon N-SIM S demo system. We acknowledge Dr. Manish Kumar Dwivedi (IISER Bhopal), Ms. Shikha Kushwaha (IISER Bhopal), Mr. Sajeev.T. K (IISER Bhopal), Mr. Satyam Sharma (IISER Bhopal) for suggestions and discussions. We also thank Rabiya Naaz for lab management.

**Supplementary video file 1:**

**On the left: Live-cell imaging showing transient contact between SNX32 vesicles and Rab11 labelled recycling endosomes:** HeLa cells co-transfected with plasmids encoding mCherry-SNX32(magenta), and GFP-Rab11(Green). Videos were captured in free run mode, without intervals in Olympus FV3000 confocal laser-scanning microscope at 37°C,5% CO_2_ with moisture control. ZDC-Z Drift compensation was used to correct focus drift during time courses. Frames were collected every 6.44 s for 09m 58s. Playback rate is 3 frames per second. The transient contact events are indicated by white arrowheads.

**On the right: Live-cell imaging showing transient contact between SNX32 vesicles and GOLPH3 labelled Golgi compartment:** HeLa cells stably expressing GFP-GOLPH3(Green) was transfected with plasmid encoding mCherry-SNX32(Magenta). Videos were captured in free run mode, without intervals in Olympus FV3000 confocal laser-scanning microscope at 37°C,5% CO_2_ with moisture control. ZDC-Z Drift compensation was used to correct focus drift during time courses. Frames were collected every 12.24 s for 09m 59s. The transient contact events are indicated by white arrowheads.

**Supplementary video file 2:**

**SNX32 colocalize with Rab11 positive recycling endosomes:**3D-reconstructed SIM movie (Nikon N-SIM S) of HeLa cell showing colocalization of mCherry-Rab11 positive recycling endosomes (Red) and GFP-SNX32 (Green). A total of 26 Z planes of 0.25µm step size was captured. For each Z plane and for each wavelength 15 images were captured (3 different angle and 5 different phases).

**Supplementary video file 3:**

**SNX32 vesicles colocalize with GOLPH3 positive Golgi compartment :**3D-reconstructed SIM movie (Nikon N-SIM S) of HeLa cell showing colocalization of GFP-GOLPH 3(Green) positive Golgi compartment and mCherry-SNX32 (Red). A total of 28 Z planes of 0.30µm step size was captured. For each Z plane and for each wavelength 15 images were captured (3 different angle and 5 different phases).

**Supplementary video file 4:**

**On the Left: SNX32, SNX4 co-traffic Transferrin:** HeLa cells co-transfected with plasmids encoding mCherry-SNX32(Magenta), and GFP-SNX4(Green) was allowed to uptake Alexa 647 labelled Transferrin (Blue) for 2 min. Videos were captured in free run mode, without intervals in Olympus FV3000 confocal laser-scanning microscope at 37°C,5% CO_2_ with moisture control. ZDC-Z Drift compensation was used to correct focus drift during time courses. Frames were collected every 9.58 s for 04m 47s. Playback rate is 2 frames per second. The co-trafficking events are indicated by white arrowheads.

**On the Right: SNX32, SNX1 co-traffic CD8-CIMPR:**

HeLa cells co-transfected with plasmids encoding mCherry-SNX1(Magenta), and GFP-SNX32(Green) and CD8-CIMPR (Blue) was processed as detailed in the materials and methods section. Videos were captured in free run mode, without intervals in Olympus FV3000 confocal laser-scanning microscope at 37°C,5% CO2 with moisture control. ZDC-Z Drift compensation was used to correct focus drift during time courses. Frames were collected every 6.520 s for 04m 49s. Playback rate is 3 frames per second. The co-trafficking events are indicated by white arrowheads.

**Supplementary video file 5:**

**SNX32 depletion hinder neurite outgrowth and network formation:** Neuro2a cells were transfected with Scrambled/SNX32/SNX6 siRNA SMART pools. Following 24hrs of transfection, the medium was replaced with MEM containing 1% fetal bovine serum supplemented with 10 μmol/L retinoic acid (RA) to induce neurite outgrowth. Frames were captured every 1 min for 48hrs using a 4× objective and a CMOS camera utilizing incubator compatible JuLI™Br Live cell analyzer. Playback rate 24 frames per sec.

**Supplementary video file 6:**

**SNX32 depletion phenocopies the neurite outgrowth defect observed in BSG downregulated condition:** Neuro2a cells were transfected with SHC002/shSNX32(#4,#6)/shBSG clones (#7,#8). Following 6hrs(shRNA) of transfection, the medium was replaced with MEM containing 1% fetal bovine serum supplemented with 10 μmol/L retinoic acid (RA) to induce neurite outgrowth. Frames were captured every 1 min for a maximum of 48hrs using a 4× objective and a CMOS camera utilizing incubator compatible JuLI™Br Live cell analyzer. Playback rate 24 frames per sec.

**Supplementary video file 7:**

shRNA resistant SNX32, shSNX32#4r over expression could rescue the neurite outgrowth defect observed in BSG downregulated condition: Neuro2a cells stably transfected with pLVXshSNX32#4r was transfected with shSNX32#4. Following 6hrs (shRNA) of transfection, the medium was replaced with MEM containing 1% fetal bovine serum supplemented with 10 ȝmol/L retinoic acid (RA) to induce neurite outgrowth. Frames were captured every 15 min for a maximum of 48hrs using a 4× objective and a CMOS camera utilizing incubator compatible JuLI™Br Live cell analyser. Playback rate 3 frames per sec.

**Supplementary video file 8:**

**SNX32 Co-traffic with BSG:** Neuro2a cells stably expressing cMyc-BSG (magenta) was co-transfected with plasmid encoding and GFP-SNX32(Green) was processed as detailed in the materials and methods section. Videos were captured in free run mode, without intervals in Olympus FV3000 confocal laser-scanning microscope at 37°C,5% CO2 with moisture control. ZDC-Z Drift compensation was used to correct focus drift during time courses. Frames were collected every 6.4 s for 04m 49s. Playback rate is 3 frames per second. The co-trafficking events are indicated by white arrowheads.

**Supplementary video file 9:**

**Surface population of pHluorin BSG is reduced in SNX32 deficit condition:** Neuro2a cells stably expressing TET inducible pHluorin BSG was transfected with Scrambled/SNX32/SNX6 siRNA SMART pools for a maximum of 72hrs. 13hrs prior to imaging the cells were induced with Doxycyclin for pHluorin BSG induction. Frames were collected every 3.3 sec for 1min. Playback rate is 3 frames per second.

**Supplementary video file 10:**

**Surface population of pHluorin BSG is reduced in SNX32 deficit condition:** Neuro2a cells stably expressing TET inducible pHluorin BSG was transfected with SHC002/SNX32(#4, #6) shRNA clones for a maximum of 42hrs. 13hrs prior to imaging the cells were induced with Doxycyclin for pHluorin BSG induction. Frames were collected every 3.3 sec for 1min. Playback rate is 3 frames per second.

## Source data legends

**Figure1-source data 1**

GBP co-immunoprecipitation of GFP /HA-tagged SNX–proteins transiently transfected in HEK293T cells showing GFP-SNX1/ GFP-SNX4/ GFP-SNX8 or GFP-SNX32 efficiently precipitating HA-SNX32, immunoblot source data of 3 biological replicates, (values represent the ratio of HA to GFP band intensity).

**Figure1-source data 2**

Co-immunoprecipitation of GFP/HA-tagged SNX–proteins transiently transfected in HEK293T cells showing GFP-SNX4 precipitating HA-SNX32ΔN, immunoblot source data of 2 biological replicates (values represent the ratio of HA to GFP band intensity).

**Figure1-source data 3**

GBP co-immunoprecipitation of GFP/HA-tagged SNX–proteins transiently transfected in HeLa cells showing difference in the amount of GFP-SNX4 precipitated HA-SNX32 mutants, immunoblot source data of 3 biological replicates (values represent the ratio of HA to GFP band intensity).

**Figure2-Source data 1**

PIP Strip membrane immunoblotted using His antibody showing preferential binding of His-SNX32ΔC to PI(3)P, PI(4)P, PI(5)P, PA (immunoblot source data of 3 biological replicates)

**Figure5-source data 1**

GBP co-immunoprecipitation of GFP tagged SNX4 and SNX32 transiently transfected in HEK293T cells showing GFP-SNX32 efficiently precipitating TfR, GBP immunoprecipitation was carried out as described in materials and methods section and immunoblotted using GFP and TfR antibody, immunoblot source data (values represent the ratio of TfR to GFP band intensity).

**Figure5-source data 2**

GBP co-immunoprecipitation of GFP tagged SNX1 and SNX32 transiently transfected in HEK293T cells showing GFP-SNX32 efficiently precipitating CIMPR, GBP immunoprecipitation was carried out as described in materials and methods section and immunoblotted using GFP and TfR antibody, immunoblot source data (values represent the ratio of CIMPR to GFP band intensity).

**Figure5-source data 3**

His affinity chromatography-based pulldown showing His-SNX32ΔC precipitating TfR from membrane enriched HeLa cell lysate fraction, His pulldown was carried out as described in material method section and immunoblotted using His and TfR antibody, immunoblot source data of 2 biological replicates (values represent the ratio of His to TfR band intensity).

**Figure5-source data 4**

His affinity chromatography-based pulldown showing His-SNX32ΔC precipitating CIMPR from membrane enriched HeLa cell lysate fraction, His pulldown was carried out as described in material method section and immunoblotted using His and TfR antibody, immunoblot source data.

**Figure5-source data 5**

Co-immunoprecipitation of GFP tagged SNX32 wild type (WT) and GFP trap of GFP-tagged SNX32 WT/ SNX32 F131D, showing both efficiently precipitating ESCPE-1 sub-unit SNX1 whereas SNX32 F131D failed to precipitate CIMPR, each transiently transfected in HEK293T cells. The elute was resolved in SDS-PAGE and immunoblotted using GFP, SNX1 and CIMPR antibody, immunoblot source data.

**Figure5-source data 6**

GBP co-immunoprecipitation of GFP tagged SNX–proteins transiently transfected in HeLa cells showing GFP-SNX32 but not GFP-SNX32 F131D efficiently pulling down TfR, GBP immunoprecipitation was carried out as described in materials and methods section and immunoblotted using GFP and TfR antibody. immunoblot source data of 3 biological replicates (values represent the ratio of TfR to GFP band intensity).

**Figure6-source data 1**

Co-immunoprecipitation of mCherry tagged SNX–proteins transiently transfected in U87MG cells showing mCherry-SNX32 but not mCherry-SNX6 efficiently pulling down Basigin (BSG), mCherry nanobody mediated pulldown was carried out as described in materials and methods section and immunoblotted using mCherry and BSG antibody, immunoblot source data of 2 biological replicates (values represent the ratio of BSG to mCherry band intensity).

**Figure6-source data 2**

Histidine (His) pulldown showing His SNX32ΔC efficientl\ pulling down BSG from membrane enriched HeLa lysate, immunoblot source data of 2 biological replicates (values represent the ratio of BSG to His band intensity).

**Figure6-source data 3**

GFP tagged SNX–proteins transiently transfected in HeLa cells showing GFP-SNX32 WT but not GFP-SNX32 F131D efficiently pulling down Basigin (BSG), GFP nanobody mediated pulldown was carried out as described in materials and methods section and immunoblotted using GFP and BSG antibody, immunoblot source data of 2 biological replicates, (values represent the ratio of BSG to GFP band intensity).

**Supplementary data Fig.S1-source data 1**

Co-immunoprecipitation of GFP /HA-tagged SNX–proteins transiently transfected in HEK293T cells showing the coprecipitation of GFP-SNX1 and HA-SNX32ΔN, GBP immunoprecipitation was carried out as described in the materials and methods section, immunoblotted using GFP and HA antibody. Immunoblot source data of 2 biological replicates.

**Supplementary data Fig.S2-source data 1**

Membrane-Cytosol fractions of HeLa cells transiently transfected with HA-SNX32ΔC or GFP-PH^OSBP^ showing PAO treatment causes delocalization of HA-SNX32ΔC and GFP-PH^OSBP^ proteins to the cytosolic fraction, S-Cytosol, P-membrane fraction, immunoblot source data (values represent the ratio of S to P fractions normalised to vinculin).

**Supplementary data Fig.S3-source data 1**

HeLa cells were transfected with Scramble/SNX32/SNX6 siRNA or SHC002, shSNX32#4, shSNX32#6 shRNA followed by cycloheximide treatment of 10 µg/ml for 6 hrs, immunoblotting was done using CIMPR or vinculin antibody (immunoblot source data of 3 biological replicates).

**Supplementary data Fig.S4-source data 1**

HeLa cells were transfected with Scramble/SNX32/SNX6 siRNA or shc002, shSNX32#4, shSNX32#6 shRNA followed by cycloheximide treatment of 10 µg/ml for 6 hrs,immunoblotting was done using TfR or vinculin antibody (immunoblot source data of 3 biological replicates).

**Supplementary data Fig.S7-source data 1**

U87MG cells were transfected with Scramble/SNX32/SNX6 siRNA or shc002, shSNX32#4, shSNX32#6 shRNA followed by cycloheximide treatment of 10 µg/ml for 6 hrs, immunoblotting was done using BSG or vinculin antibody (immunoblot source data of 3 biological replicates).

---

**Supplementary data Fig.S 1.**
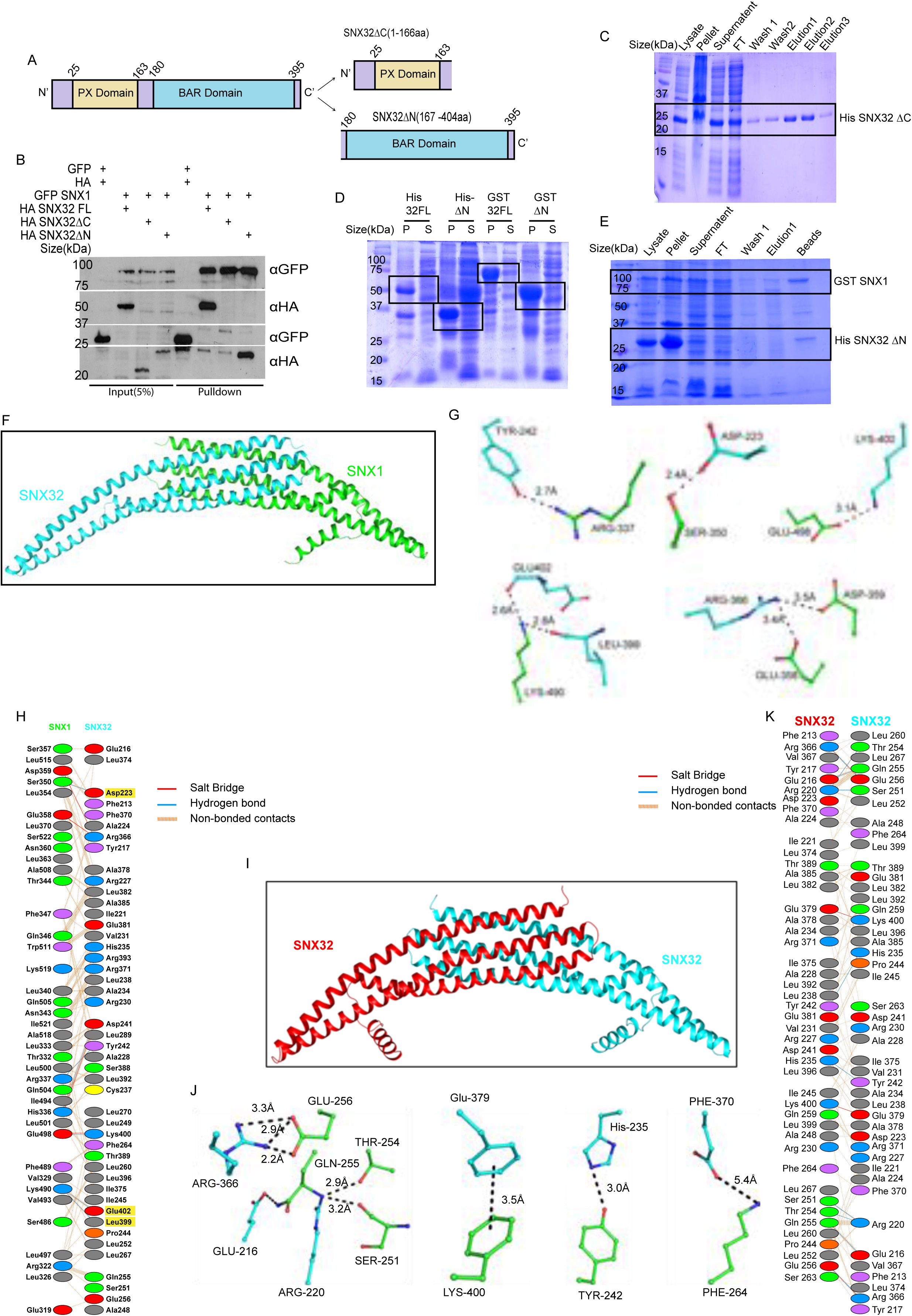
SNX32 undergoes BAR domain mediated association with SNX1: A) The domain architecture indicating the N- and C-terminal endpoints of SNX32FL, SNX32ΔC and SNX32ΔN. B) Co-immunoprecipitation of GFP /HA-tagged SNX–proteins transiently transfected in HEK293T cells showing the coprecipitation of GFP-SNX1 and HA-SNX32ΔN, GBP immunoprecipitation was carried out as described in the materials and methods section, immunoblotted using GFP and HA antibody. C)Coomassie blue-stained SDS-PAGE gel of affinity purification profile of His SNX32ΔC, samples representing each step of purification, FT: Flow-through. D) Coomassie blue-stained SDS-PAGE gel of His-SNX32 FL, His-SNX32ΔC, GSTSNX32FL, GST-SNX32 ΔC after induction, P: pellet and S: supernatant fractions. E) Coomassie blue-stained SDS-PAGE gel of affinity co-purification profile of GST-SNX1/His-SNX32ΔN, samples representing each step of purification, FT: Flow-through. F) homology model of SNX32 BAR domain(cyan) in complex with SNX1 BAR domain(green). G) Interacting amino acid residues present in the dimeric interface of SNX32(cyan)-SNX1(green). H) Snap shots of polar interactions across the dimeric interface. I)homology model of SNX32 BAR homodimer J) Interacting amino acid residues present in the dimeric interface of SNX32(cyan)-SN32(red). K) Snap shots of polar interactions across the dimeric interface

**Supplementary data Fig.S 2.**
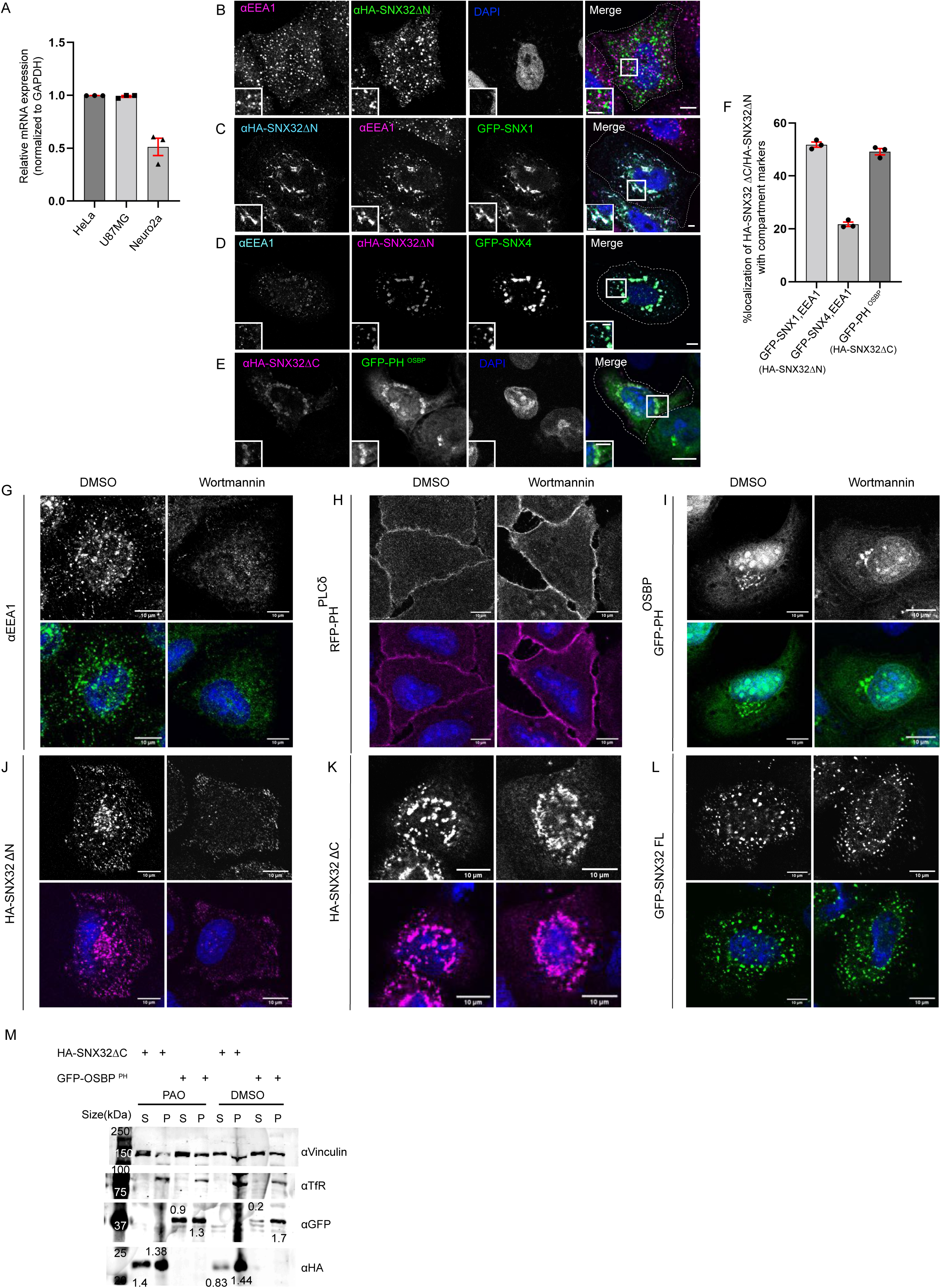
BAR domain of SNX32 undergoes protein-protein interactions and assist in the early endosomal localization of SNX32: A) Relative amount of SNX32 transcripts analysed by quantitative PCR in different cell lines such as HeLa, U87MG and Neuro2a normalised with respect to Gapdh. B-D) Localization of HA SNX32ΔN with B) early endosomal marker EEA1 C) GFP SNX1, EEA1 G) GFP-SNX4, EEA1: Scale bar 10µm, inset 5µm (magnified regions are shown as insets). E) HA SNX32ΔC colocalizes with GFP PH^OSBP^, protein module showing preferential association with PI(4)P, Scale bar 10µm, inset 5µm (magnified regions are shown as insets). F) Quantifications showing percentage cRlRcali]aWiRQ Rf HA SNX32ΔN/ HA SNX32ΔC with respective compartment markers in HeLa cells, data represent mean ±SEM (N=3, n≥60cells per independent experiments). G-L) Wortmannin/DMSO treatment in HeLa cells overexpressing G) endogenous EEA1, H) RFP-PH^PLCδ^, I) GFP-PH^OSBP^, J) HA-SNX32ΔN, K) HA-SNX32ΔC, L) HA-SNX32FL, Scale bar 10µm. M)Membrane-Cytosol fractions of HeLa cells transiently transfected with HA-SNX32ΔC or GFP-PH^OSBP^ showing PAO treatment causes delocalization of HA-SNX32ΔC and GFP-PH^OSBP^ proteins to the cytosolic fraction, S-Cytosol, P-membrane fraction (representative immunoblot out of 3 biological replicates, values represent the ratio of S to P fractions normalised to vinculin).

**Supplementary data Fig.S 3.**
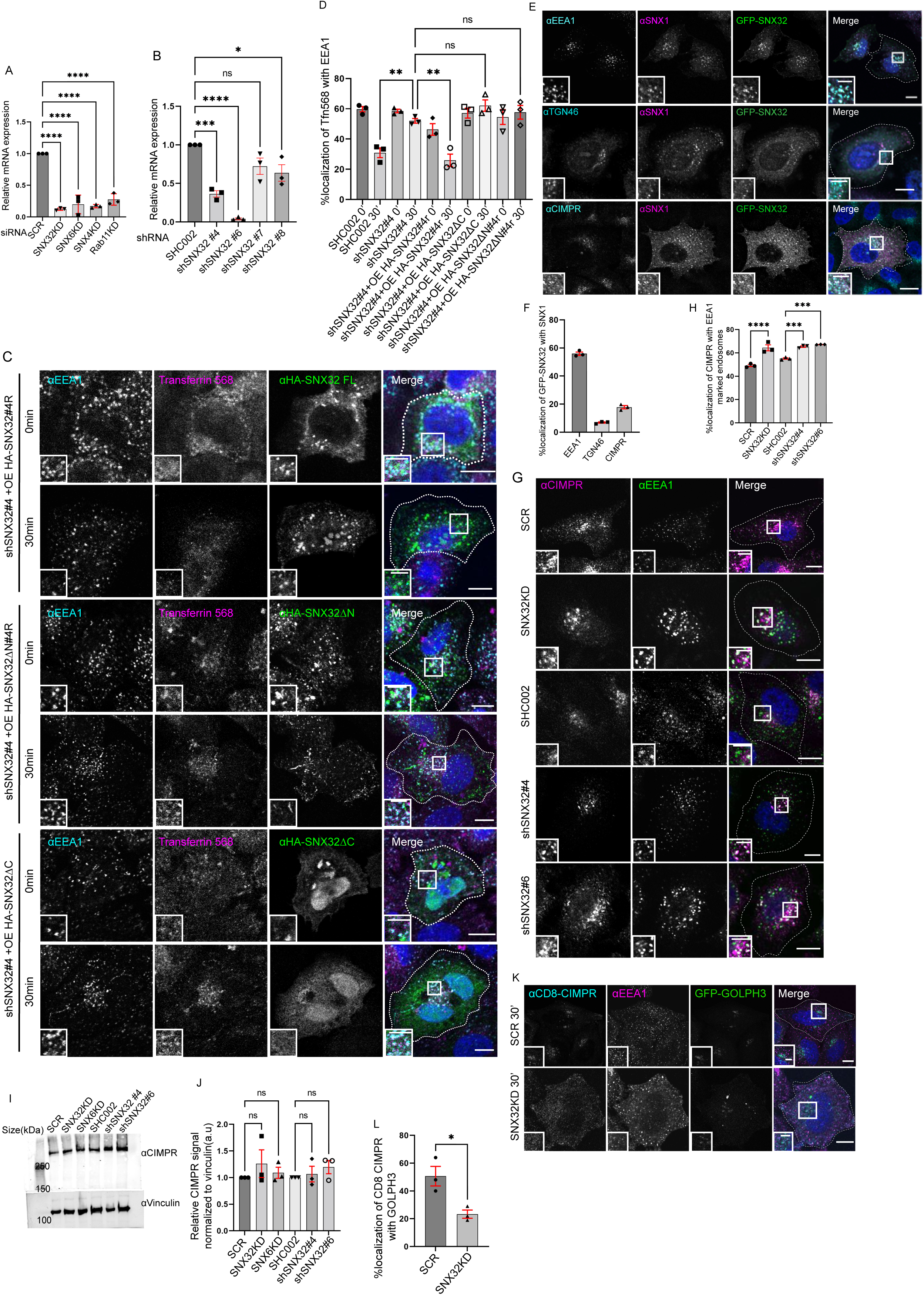
SNX32 interacts with and regulate the trafficking of CIMPR in HeLa cells: A) Analysis of the relative gene expression levels of SNX32, SNX6, SNX4 and Rab11 by qRT-PCR in HeLa cells depleted for SNX32, SNX6, SNX4 and Rab11 respectively. Values of control were arbitrarily set as 1 against which experimental data were normalized. Gapdh was used as internal control, data represent mean ±SEM (N=3). B) Analysis of the relative gene expression levels of SNX32 by qRT-PCR in HeLa cells depleted for SNX32 utilising different shRNA clones. Values of control were arbitrarily set as 1 against which experimental data were normalized. Gapdh was used as internal control, data represent mean ±SEM (N=3). C) Followed by shSNX32#4 mediated gene down regulation of SNX32 HA-SNX32#4r/ HA-SNX32ΔN#4U RU HA-SNX32ΔC was over expressed, the transferrin (Alexa Fluor 568 conjugated) Pulse-Chase experiment was carried out as described in materials and method section, the cells were fixed at specified timepoints, immunostained using early endosomal marker EEA1, DAPI was used to stain nucleus, Scale 10µm, inset 5µm (magnified regions are shown as insets).D) Quantification of percentage localization of transferrin (Alexa Fluor 568 conjugated) with EEA1 at corresponding time points, data represent mean ±SEM (N=3, n≥15 random frames per independent experiments), P value <0.0001, Ordinary one-way ANOVA, Šídák’s multiple comparison test (** P < 0.01, ns-non significant). E-F) SNX32 and SNX1 colocalizes with (E) early endosome marker EEA1, Trans Golgi Network marker TGN46, CIMPR, Scale bar 10µm, inset 5µm (magnified regions are shown as insets), F) Percentage localization of SNX32-SNX1 heterodimer with EEA1, TGN46, and CIMPR, data represent mean ±SEM (N=3, n≥60cells per independent experiments). G)Followed by SMARTpool mediated gene downregulation of Scrambled (SCR) / SNX32 or shRNA mediated gene downregulation of scrambled (SHC002) / SNX32 shRNA clones(shSNX32#4,shSNX32#6) HeLa cells where immunostained using CIMPR, EEA1 and nuclei were counterstained with DAPI (blue), Scale 10µm, inset 5µm (magnified regions are shown as insets), H) Quantification of percentage localization of CIMPR with EEA1, data represent mean ±SEM (N=3, n≥60cells per independent experiments), P value <0.0001(*** P < 0.001,****P < 0.0001), One way ANOVA, Šídák’s multiple comparison test. I)HeLa cells were transfected with Scramble/SNX32/SNX6 siRNA or SHC002, shSNX32#4, shSNX32#6 shRNA followed by cycloheximide treatment of 10 µg/ml for 6 hrs, immunoblotting was done using CIMPR or vinculin antibody (representative immunoblot out of 3 biological replicates). J) Quantification of relative CIMPR signal normalized to vinculin (loading control), data represent mean ±SEM (N=3) P value 0.7104, One-way ANOVA Šídák’s multiple comparisons test. K) Followed by SMARTpool mediated gene downregulation of Scrambled (SCR)/SNX32 in HeLa cells stably co-expressing GFP-GOLPH3 and CD8-CIMPR (CD8-CIMPR is a chimeric protein in which the cytoplasmic tail of CD8, cell surface protein, is replaced with the cytoplasmic tail of CIMPR, enabling CIMPR to shuttle from the plasma membrane to TGN), CD8 antibody uptake experiment was carried out as described in materials and method section, the cells were fixed at 30min timepoint, immunostained using EEA1 and nuclei were counterstained with DAPI (blue), Scale 10µm, inset 5µm (magnified regions are shown as insets). L)Quantification of percentage localization of CD8-CIMPR with GFP GOLPH3, data represent mean ±SEM (N=3, n≥60cells per independent experiments), P value 0.0233, Unpaired t test Two-tailed (* P < 0.05).

**Supplementary data Fig. S4.**
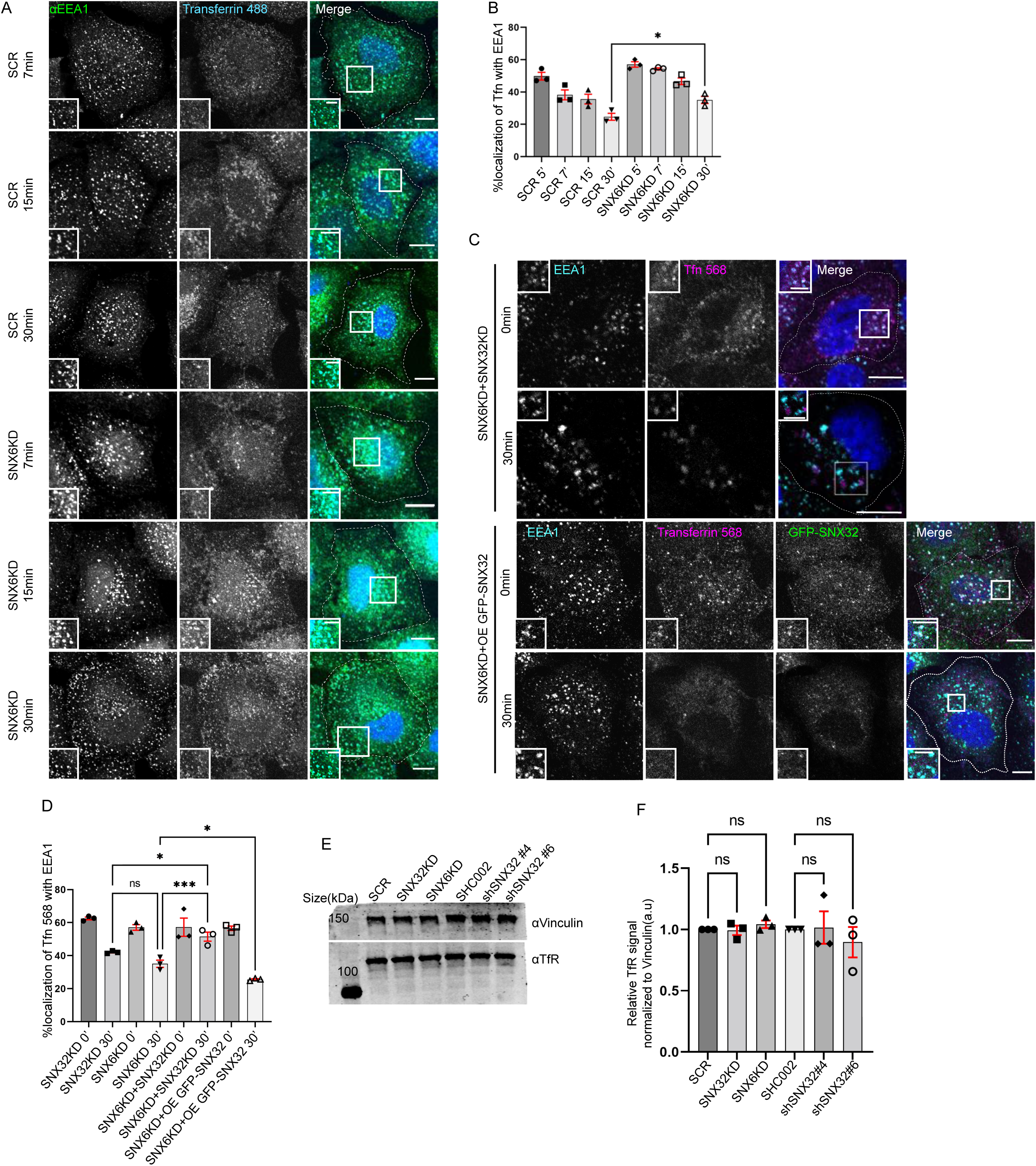
SNX6 and SNX32 show an overlapping role in regulating the trafficking of transferrin in HeLa cells: A) Followed by SMARTpool mediated gene downregulation of Scrambled (SCR) or SNX6, transferrin (Alexa Fluor 488 conjugated) Pulse-Chase experiment was carried out as described in materials and method section, the cells were fixed at specified time points, immunostained using EEA1 and Nuclei were counterstained with nuclei were counterstained with DAPI (blue) (blue), Scale 10µm, inset 5µm (magnified regions are shown as insets). B) Quantification of percentage localization of transferrin (Alexa Fluor 488 conjugated) with EEA1 at corresponding time points, data represent mean ±SEM (N=3, n≥60cells per independent experiments), P value <0.0273 Unpaired t test Two-tailed (* P < 0.05).C) Followed by SMARTpool mediated gene down regulation of SNX6 and SNX32 or SNX6KD and over expression of GFP-SNX32, the transferrin (Alexa Fluor 568 conjugated) Pulse-Chase experiment was carried out as described in materials and method section, the cells were fixed at specified timepoints, immunostained using early endosomal marker EEA1, DAPI was used to stain nucleus, Scale 10µm, inset 5µm (magnified regions are shown as insets). D) Quantification of percentage localization of transferrin (Alexa Fluor 568 conjugated) with EEA1 at corresponding time points, data represent mean ±SEM (N=3, n≥15 random frames per independent experiments) P value <0.0001 one-way ANOVA Šídák’s multiple comparisons test (* P < 0.05, *** P < 0.001, ns-nonsignificant). E) HeLa cells were transfected with Scramble/SNX32/SNX6 siRNA or shc002, shSNX32#4, shSNX32#6 shRNA followed by cycloheximide treatment of 10 µg/ml for 6 hrs, immunoblotting was done using TfR or vinculin antibody (representative immunoblot out of 3 biological replicates). F) Quantification of relative TfR signal normalized to vinculin (loading control), data represent mean ±SEM (N=3) P value 0.8254, One-way ANOVA Šídák’s multiple comparisons test (ns-nonsignificant).

**Supplementary data Fig. S5.**
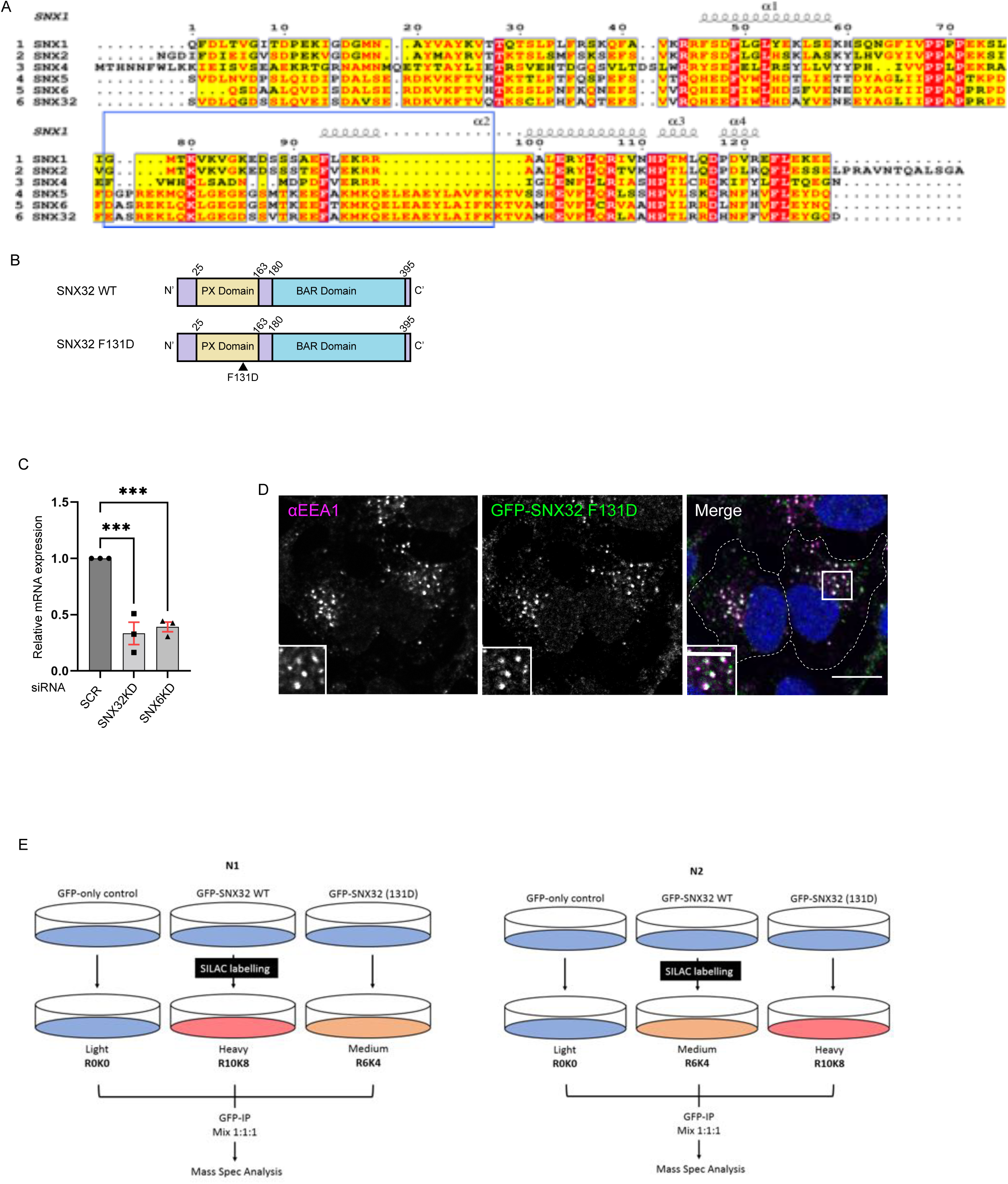
Mapping the interactome of SNX32, utilising the conserved F131 residue: A) Alignment of the PX domain of SNX1, SNX2, SNX4, SNX5, SNX6, SNX32. The blue box represents the α’ and α’’ helices that compose the unique helix-turn-helix extension, the black arrow points to F131 residue of SNX32. B) The domain architecture of SNX32 indicating the N- and C-terminal endpoints of SNX32FL, SNX32ΔC and SNX32 F131D. C) Quantification of relative mRNA expression of SNX32 and SNX6 in Neuro2a cells treated with respective SMART Pool siRNAs, Values of control were arbitrarily set as 1 against which experimental data were normalized. Gapdh was used as internal control, (N=3, values are means ± SEM), P value 0.0005 (*** P < 0.001), Ordinary one-way ANOVA Dunnett’s multiple comparisons tests. D) SHSY5Y cells showing colocalization of GFP tagged SNX32 F131D with EEA1 harbouring endosomes, Scale 10µm, inset 5µm (magnified regions are shown as insets). E) Schematic representation of the SILAC methodology employed to obtain the differential proteomic dataset.

**Supplementary data Fig. S6.**
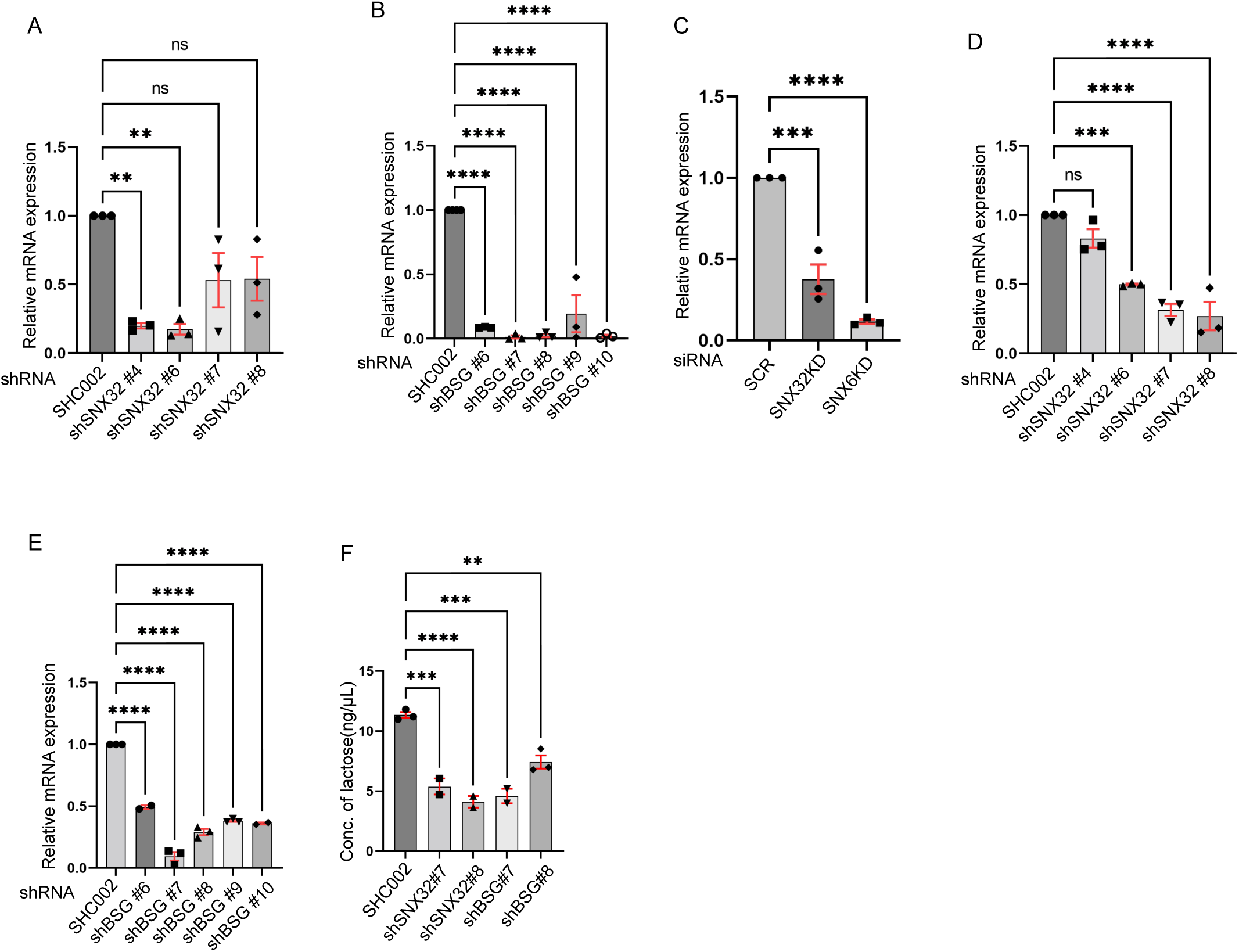
Efficiency of target gene depletion under siRNA and shRNA transfection, estimated based on mRNA expression levels: Relative mRNA expression of Neuro2a cells transfected with A) SNX32 shRNA clones, P value 0.0030 (N=3, values are means ± SEM, ** P < 0.01, ns-nonsignificant), Ordinary one-way ANOVA Dunnett’s multiple comparisons tests. B) BSG shRNA clones, P value <0.0001, SMART Pool siRNAs, (N=3, values are means ± SEM, **** P < 0.0001), Ordinary one-way ANOVA Dunnett’s multiple comparisons tests. C) Quantification of relative mRNA expression of SNX32 and SNX6 in U87MG cells treated with respective SMART Pool siRNAs, (N=3), P value <0.0001 (*** P < 0.001, **** P < 0.0001), Ordinary one-way ANOVA Dunnett’s multiple comparisons tests. D-E) Relative mRNA expression of U87MG cells transfected with D) SNX32 shRNA clones, P value <0.0001, E) BSG shRNA clones, P value <0.0001, SMART Pool siRNAs, (N=3, values are means ± SEM, *** P < 0.001, **** P < 0.0001, ns-nonsignificant), Ordinary one-way ANOVA Dunnett’s multiple comparisons tests. F) Quantification of concentration of lactate in the culture supernatant of U87MG cells transfected with Scrambled (SHC002)/SNX32 shRNA clones (shSNX32#4, shSNX32#6)/ BSG shRNA clones (shBSG#7, shBSG#8), N=2, P value <0.0001(** P < 0.01, *** P < 0.001, **** P < 0.0001), Ordinary one-way ANOVA Dunnett’s multiple comparisons test.

**Supplementary data Fig. S7.**
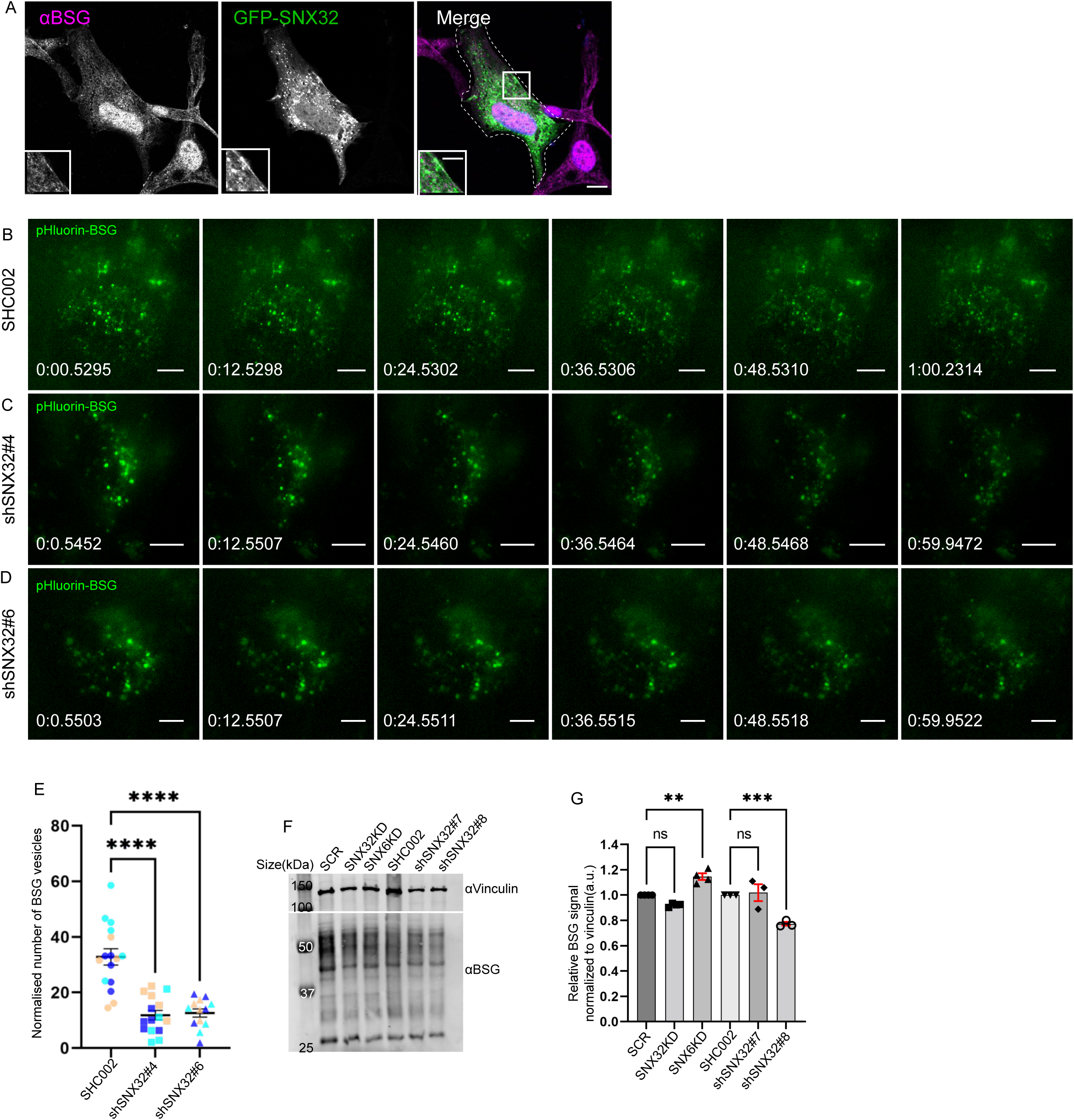
SNX32 colocalize and regulates the surface population of BSG: A-B) B-D) Snapshots from live TIRF microscopic imaging of Neuro2a cells stably expressing pHluorin BSG transfected with B) SHC002 or SNX32 shRNA clones C) shSNX32#4, D) shSNX32#6 followed by doxycycline treatment for pHluorin BSG induction, Scale 10µm. E) Quantification of the surface population of the number of BSG vesicles (N=3, n≥8cells per independent experiments), P value <0.0001(**** P < 0.0001), Ordinary one-way ANOVA Dunnett’s multiple comparisons tests. F) U87MG cells were transfected with Scramble/SNX32/SNX6 siRNA or shc002, shSNX32#4, shSNX32#6 shRNA followed by cycloheximide treatment of 10 µg/ml for 6 hrs, immunoblotting was done using BSG or vinculin antibody (representative immunoblot out of 3 biological replicates). G) Quantification of relative BSG signal normalized to vinculin (loading control), data represent mean ±SEM (N=3) P value <0.0001, (** P < 0.01, *** P < 0.001) One-way ANOVA Šídák’s multiple comparisons test.

